# Mouse scalp development requires Rac1 and SRF for the maintenance of mechanosensing mesenchyme

**DOI:** 10.1101/2025.02.11.637680

**Authors:** Bharath H. Rathnakar, Alex Rackley, Hae Ryong Kwon, William L. Berry, Lorin E. Olson

## Abstract

Regulation of essential cellular responses like proliferation, migration, and differentiation is crucial for normal development. Rac1, a ubiquitously expressed small GTPase, executes these responses under the regulation of guanine nucleotide exchange factors (GEFs) and GTPase activating proteins (GTPases). Mutations in specific GEFs (DOCK6) and GTPases (AHGAP31) that regulate Rac1 are associated with Adams-Oliver syndrome (AOS), a developmental syndrome characterized by congenital scalp defects and limb truncations. Genetic ablation of *Rac1* in the mouse embryonic limb ectoderm results in limb truncation. However, the etiology of Rac1-associated cranial defects is unknown. To investigate the origin and nature of cranial defects, we used a mesenchymal Cre line (*Pdgfra-Cre*) to delete *Rac1* in cranial mesenchyme. *Rac1*-KO mice died perinatally and lacked the apical portion of the calvarium and overlying dermis, resembling cranial defects seen in severe cases of AOS. In control embryos, α-smooth muscle actin (αSMA) expression was spatially restricted to the apical mesenchyme, suggesting a mechanical interaction between the growing brain and the overlying mesenchyme. In *Rac1*-KO embryos there was reduced proliferation of apical mesenchyme, and reduced expression of αSMA and its regulator, serum response factor (SRF). Remarkably, *Srf*-KO mice generated with *Pdgfra-Cre* recapitulated the cranial phenotype observed in *Rac1-*KO mice. Together, these data suggest a model where Rac1 and SRF are critical to maintaining apical fibroblasts in a mechano-sensitive and proliferative state needed to complete cranial development.

## INTRODUCTION

Rho GTPases are molecular switches that control many aspects of cell biology by cycling between two conformational states. An active conformation (GTP-bound) engages with effector proteins while GTPase-mediated hydrolysis returns them to an inactive conformation (GDP-bound). The conformational change in Rho GTPases is controlled by two classes of regulators with opposing functions, guanine nucleotide exchange factors (GEFs) and GTPase-activating proteins (GTPases) (Jaffe and Hall 2005; Ridley 2011; Duquette and Lamarche-Vane 2014; Hodge and Ridley 2016). RAS-related C3 Botulinum Toxin Substrate 1 (RAC1) is a member of the Rho family of GTPases, including RAC2, RAC3, and RhoG (Aspenstrom et al. 2004; Boureux et al. 2007; Reijnders et al. 2017). Rac1 is ubiquitously expressed and transduces signals received from extracellular stimuli (growth factors/receptor tyrosine kinases, G-protein coupled receptors and integrins) via activated GEFs (DOCK6) through its effectors (WAVE, IQGAP, PAK and MLK2/3) (Ridley et al. 1992; Sugihara et al. 1998; Bosco et al. 2009) to induce lamellipodia/membrane ruffles to modulate cell growth, cytoskeleton organization, reactive oxygen species (ROS) generation, and mesenchymal-like migration (Aspenstrom et al. 2004; Bosco et al. 2009; Saci et al. 2011; Wojnacki et al. 2014). Mutations in Rac1 or its regulators can lead to developmental defects.

Adams-Oliver Syndrome (AOS) is a multiple congenital anomaly with an incidence ratio of 1 in 225,000 births (Saeidi and Ehsanipoor 2017). It was first reported in 1945 with 8 cases in one family (ADAMS and OLIVER 1945; Whitley and Gorlin 1991) . It is characterized by aplasia cutis congenita (ACC) with calvaria defects and terminal transverse limb defects (TTLD) (Southgate et al. 2015). AOS individuals present defects of the apical head including alopecia, congenital absence of skin or scalp on the skull vertex and scarred atrophic plaque (Southgate et al. 201 ; Sukalo et al. 2015; Rashid et al. 2022). In severe cases, calvaria agenesis exposes the dura, leading to herniation, and bleeding of the brain (Wehrens et al. 2020). AOS individuals also present terminal transverse limb defects (TTLD), commonly in the distal limb with hypoplastic digits, syndactyly, or complete absence of a finger, toe, or distal limbs (Seo et al. 2010; Bakry et al. 2012; Saeidi and Ehsanipoor 2017; Rashid et al. 2022). To date, AOS has been linked with several genes. Mutations in ARHGAP31 (MIM#610911) (Southgate et al. 2011), RBPJ (MIM#147183) (Hassed et al. 2012) DLL4 (MIM#616589) (Meester et al. 2015), and NOTCH1 (MIM#190198) (Stittrich et al. 2014) are inherited as a dominant trait. Mutations in DOCK6 (MIM#614194) (Shaheen et al. 2011) and EOGT (MIM#614789) (Shaheen et al. 2013) exhibit autosomal-recessive transmission. Among these genes, ARHGAP31 and DOCK6 are Rho GTPase regulators primarily involved in controlling the activity of Rac1 and a related GTPase, Cdc42 (Lamarche-Vane and Hall 1998; Cote and Vuori 2002; Jenna et al. 2002; Cote and Vuori 2006; Laurin and Cote 2014). ARHGAP31 facilitates the exchange of GTP-GDP, thereby rendering the conformation of Rac1 to its inactive state. On the other hand, DOCK6 is a critical GEF that facilitates GDP-GTP exchange to activate Rac1. Gain of function mutation in ARHGAP31 and loss of function mutation in DOCK6 lead to sustained inactivation of Rac1, which affects actin remodeling consistent with the established role of Rho GTPases in organizing the actin network (Shaheen et al. 201 ; Southgate et al. 2015; Sukalo et al. 2015). This hints at the possibility that deregulated Rac1 activity could be the underlying cause of AOS. So far, AOS mutations in mouse ARHGAP31 and DOCK6 have not been described.

In mouse studies, global deletion of *Rac1* is embryonic lethal as it is required during gastrulation (Sugihara et al. 1998). Intriguingly, conditional deletion of *Rac1* using *Msx2-Cre,* which targets the apical ectodermal ridge (AER) but not the mesenchyme of the mouse embryonic limb bud, resulted in the lack of hindlimb structures and truncation of forelimb at various levels (Wu et al. 2008). Thus, *Rac1* deletion from the limb ectoderm recapitulated the limb truncation seen in AOS individuals harboring mutations in ARHGAP31 and DOCK6. In contrast, conditional deletion of *Rac1* using *Prrx1-Cre*, which targets the limb mesenchyme, generated mice with shorter ulnae and tibiae but no truncations. Webbing of interdigital skin was observed in these mice representing syndactyly of soft tissue (Suzuki et al. 2009). These reports indicate the close similarities of limb truncation phenotype between AOS patients and in mice with ectodermal deletion of *Rac1*, but not in mice with mesenchymal deletion of *Rac1*. This suggests that regulation of Rac1 in limb ectoderm by DOCK6 and ARHGAP31 may be important for limb development. Nevertheless, the calvaria phenotype seen in AOS individuals has not been recapitulated in mice.

We performed genetic deletion of *Rac1* using *Pdgfra-Cre* to target cranial mesenchyme. Our results show that *Pdgfra-Cre;Rac1^Flox/Flo^*^x^ mice recapitulate the cranial phenotype seen in severe AOS individuals. The calvaria defects were localized to the apical portion of the frontal, parietal, and interparietal bones. Analysis revealed hypoplasia of apical fibroblasts with reduced expression of serum response factor (SRF) and α-smooth muscle actin (αSMA). Intriguingly, the *Pdgfra-Cre;Srf^Flox/Flox^* phenotype was very similar to that of the *Rac1* mutation. These results demonstrate *Rac1*-regulated calvaria development in mice, the failure of which corresponds to the phenotype observed in AOS individuals. This suggests a population of PDGFRα^+^ mechano-sensitive apical fibroblasts that express αSMA in response to the mechanical stress produced by the growing brain confronting the overlaying cranial mesenchyme. This response is critical in the spatiotemporal emergence of mechano-sensitive apical fibroblasts at the skull vertex where the ability of Rac1 and SRF to relay mechanotransduction signaling is important for the development of the apical skull and dermis.

## RESULTS

### Deletion of *Rac1* in the PDGFRα^+^ mesenchymal cells causes scalp and calvaria defects with perinatal lethality

Global deletion of *Rac1* is embryonic lethal as it is required to form germ layers during gastrulation (Sugihara et al. 1998). Therefore, we used a mesenchymal Cre line, *Pdgfr*a*-Cre^Tg^* (Berry and Rodeheffer 2013; Rivera-Gonzalez et al. 2016), to knock out *Rac1* in PDGFRα^+^ cells. *Pdgfra-Cre^Tg^* mice were crossed with *Rac1^Flox/Flox^ (Rac1^tm1Djk/^J)* mice (Glogauer et al. 2003) as represented in Figure 1A to generate mutant *Pdgfra-Cre;Rac1^Flox/Flox^* embryos (hereafter *Rac1-*KO) and *Pdgfra-Cre Rac1^Flox/+^ or Rac1^Flox/Flox^* control embryos (hereafter *Rac1*-Wt)*. Rac1*-KO mice were born with the scalp and apex of the skull missing, leading to herniation of the brain (Fig. 1B). Although the *Rac1-*KO pups were born alive, they were significantly below the expected Mendelian ratio with 8 mutants out of 76 pups at postnatal day 1 (P1) when 19 mutants were expected (Fig. 1C). There were 11 mutants out of 73 pups at E18.5 when 18 mutants were expected (Suppl. Fig. 1A). Most *Rac1*-KO pups did not survive beyond P1, except for one that survived until P3 (Suppl.Fig.1B), demonstrating perinatal lethality associated with *Rac1*-KO in mice. As shown below, western blot and qPCR confirmed deletion of Rac1 in cultured dermal fibroblasts (Fig. 5A-C).

**Figure 1:**
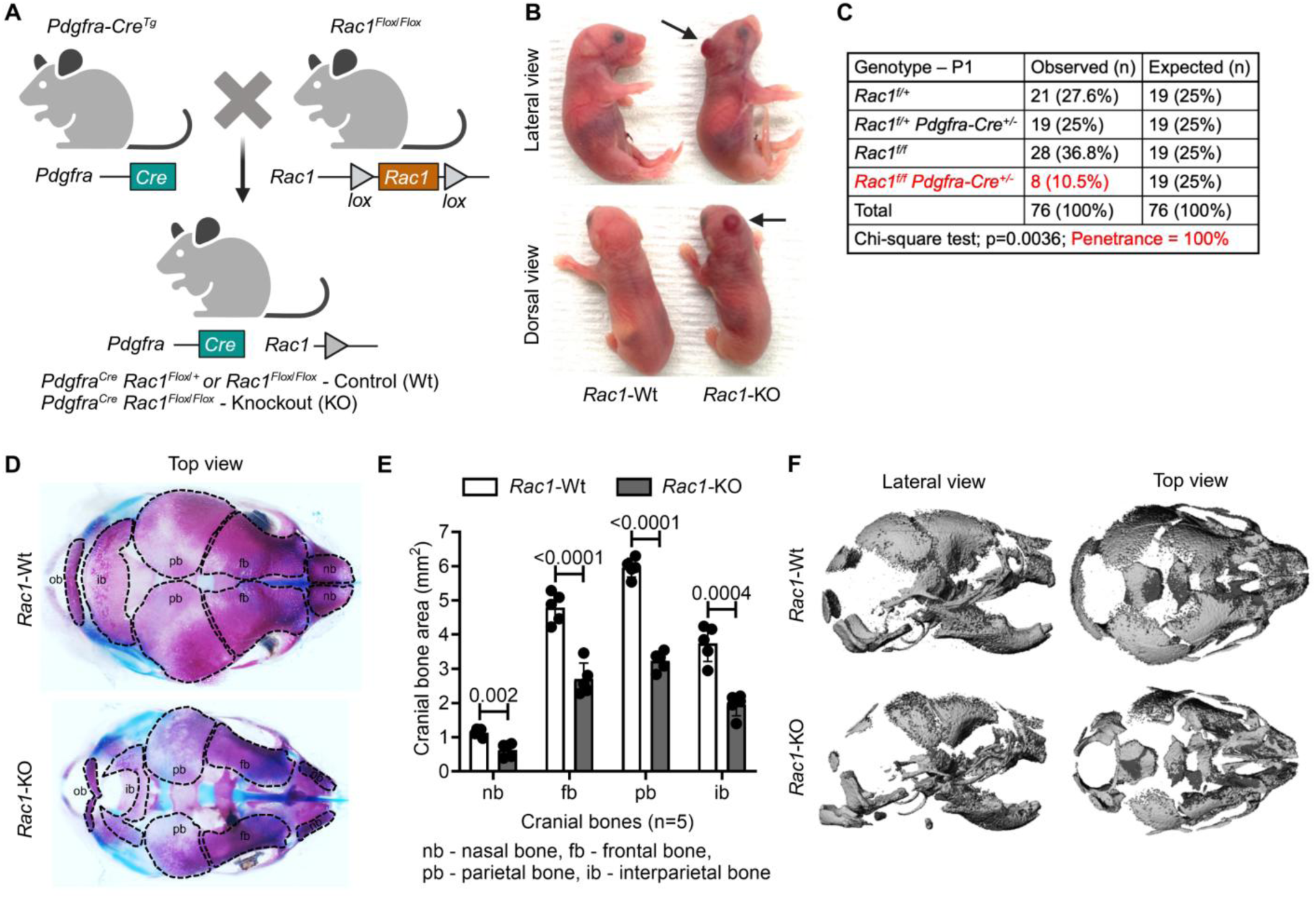
Mesenchymal deletion of *Rac1* causes scalp and calvaria defects and perinatal lethality. (A) Conditional knockout strategy for *Pdgfra^Cre^ Rac1^Flox/Flox^*. (B) Lateral and dorsal view of *Rac1*-Wt and -KO P1 pups. Arrow indicates the scalp defect (n=8/8). (C) Genotypes of pups at P1, two-tailed Chi-square test of observed genotypes. 100% of *Pdgfra^Cre^ Rac1^Flox/Flox^* mice exhibited defects shown in Fig.1B. (D) Skeletal preparation of P1 skull with alizarin red and alcian blue staining (n=5). Dotted lines outline the individual bones of the calvarium. (E) Quantification of cranial bones (n=5). Data represented as mean ± SD, Unpaired t-test with Welch’s correction. (F) Micro CT surface rendering of skulls at P1 showing loss of calvaria bone volume (n=1).

We performed skeletal preparations to characterize the calvaria phenotype at P1. The *Rac1*-KO skulls displayed dysplasia of the nasal bones, the frontal, parietal, interparietal, and occipital bone at the midline (Fig. 1D). Quantification revealed significantly reduced size of the nasal bones, frontal bones, parietal bones, and intraparietal bones of *Rac1-*KO compared to *Rac1-*Wt littermates (Fig. 1E). The occipital bone was also reduced in *Rac1-*KO (Fig. 1D). A Micro-CT rendering of the skull indicated dramatic loss of mineralized calvaria bone in *Rac1-*KO (Fig. 1F). Nevertheless, the skeleton of P1 *Rac1*-KO mice displayed no additional defects, except for the overall size of the skeleton being slightly reduced in *Rac1-*KO compared to *Rac1-*Wt (Suppl. Fig.1C). These data demonstrate the significance of Rac1 in PDGFRα^+^ mesenchymal cells during mouse calvaria genesis. Intriguingly, the *Rac1-*KO phenotype resembles skull and scalp defects seen in severe cases of AOS (ADAMS and OLIVER 1945; Sukalo et al. 2015; Wehrens et al. 2020). However, AOS-like limb truncation was not recapitulated in *Rac1*-KO, suggesting different mechanisms for limb and apical head development.

### *Pdgfra-Cre^Tg^* is expressed in cephalic mesenchyme at E11.5

Despite *Rac1* being ubiquitously expressed and *Pdgfra* being broadly expressed in embryonic mesenchyme, the phenotype was localized to the apical region of the head. PDGFRα is known to be expressed in neural crest cells that give rise to many craniofacial structures (Tallquist and Soriano 2003; He and Soriano 2013), but no studies have reported the spatiotemporal expression of *Pdgfra-Cre* during development. To address this question, we crossed *Pdgfra-Cre^Tg^* mice with *ROSA26^Ai4^* mice (Madisen et al. 2010) (Suppl. Fig. 2A). At E11.5, the highest Tomato expression was observed in the cephalic mesenchyme (Suppl. Fig. 2B-C). At E13.5, Tomato was expressed in the apical and basolateral cephalic mesenchyme that gives rise to meninges, dermis, and bone (Dasgupta and Jeong 2019). Tomato was expressed in limb mesenchyme only at E13.5 (Suppl. Fig. 2B), indicating significantly delayed expression of *Pdgfra-Cre^Tg^* compared to the endogenous *Pdgfra* expression at 7.5-8.5 (Orr-Urtreger et al. 1992). This could partly explain why the major phenotype was localized to the apical head (Fig. 1B, D-F). Most likely *Pdgfra-Cre^Tg^* activity starts at E10.5, as we first saw weak expression of Tomato in the head at this time. These results confirm Cre activity in PDGFRα^+^ cephalic mesenchyme in agreement with the defects seen in *Rac1*-KO mice.

### Ontogeny of the *Rac1*-KO cranial phenotype

To determine the earliest occurrence of a cranial defect in *Rac1-*KO embryos, timed pregnancies were conducted. The earliest phenotype was observed at E11.5 as a slight enlargement of the caudal head (Fig. 2A), followed by a slight change in the shape of the caudal head and a distinct blebbing on the nose at E12.5 (Fig. 2A). At E13.5 and E14.5, the caudal deformity enlarged into a large blebbing, and the nasal bleb became a hematoma (Fig. 2A). However, the nasal blebbing and hematoma were resolved by E18.5 and only the caudal defect in the apical head remained externally obvious (Fig. 2A). Interestingly, one of the mutants among the 16 *Rac1*-KO embryos observed at E14.5 exhibited mid-facial cleft (Suppl. Fig. 3A). Previously, *Thomas et al*., reported a phenotype with a severe mid-facial cleft at E12.5 when *Rac1* was deleted in neural crest cells using *Wnt1-Cre* (Thomas et al. 2010). PDGFRα is expressed in neural crest cells (Soriano 1997; Tallquist and Soriano 2003; He and Soriano 2013; Mo et al. 2020), but *Pdgfra-Cre* targets them at a later time than *Wnt1-Cre*, thereby contributing to the low penetrance of mid-facial cleft when used to generate *Rac1*-KO.

**Figure 2:**
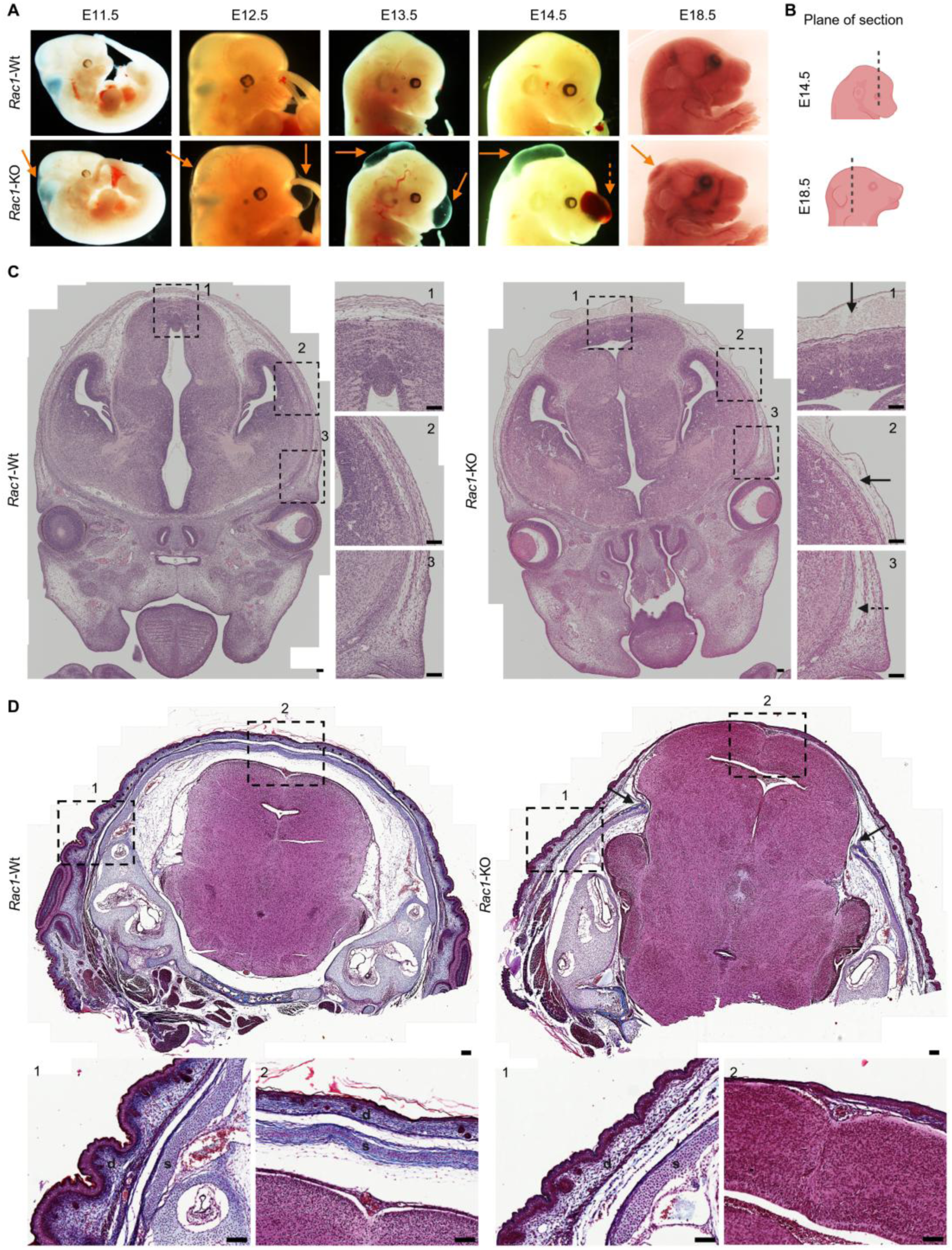
Mesenchymal *Rac1*-KO causes aplasia of the apical dermis and calvaria. (A) Lateral view of *Rac1*-Wt and -KO embryos from E11.5 to E18.5. The orange arrows indicate blebbing/edema, hematoma (dotted arrow), and scalp defect (n=2-19). (B) Graphical representation of coronal plane of section for E14.5, and E18.5 mouse embryos shown in C-D. (C) H&E staining of coronal sections of *Rac1*-Wt and -KO E14.5 embryos. Dotted box=insets, dotted arrow=hypoplasia of basolateral mesenchyme, arrow=blebbing/edema of intermediate and apical mesenchyme. The insets are represented as 1=apical, 2=intermediate, and 3=basolateral region of the head (n=3). Scale=50μm. (D) Trichrome staining of coronal sections of *Rac1*-Wt and -KO E18.5 head. Arrows in *Rac1*-KO indicate the apical limits of the intact calvaria, with the brain protruding through the gap and covered by a thin layer of the rudimentary dermis. Scale=50μm. The dotted box = Inset. The bottom panel represents the inset 1 = basolateral, 2 = apical, d = dermis, and s = skull (n=2). Scale=50μm.

Sagittal sections at E14.5 revealed edema of the caudal mesenchyme and nasal hematoma in *Rac1-* KO (Suppl. Fig. 3B). Coronal sections revealed hypoplasia of the *Rac1-*KO dermis lateral to the pre-osteogenic layer (Fig. 2C, inset 3), which became edematous in the intermediate region towards the apex (Fig. 2C, inset 2). Edema was most severe in the mesenchyme at the apex of the head (Fig. 2C, inset 1). Brain shape was altered at E14.5 and E18.5 (Fig. 2C-D), but we did not explore this in detail because we considered brain changes to be secondary to cranial defects arising from mesenchymal cells. Coronal sections of *Rac1*-KO at E18.5 (Fig. 2D) revealed a lack of calvaria and only a rudimentary apical dermis covering the brain (Fig. 2D, inset 2), while basolateral regions of calvaria and dermis were intact (Fig. 2D, inset 1). In summary, the cranial defect began at E11.5-E14.5 as mesenchymal hypoplasia and edema over a large portion of the head and dorsum of the embryo, plus nasal hematoma. However, by E18.5 the nasal hematoma was resolved and only the calvaria and apical dermis were severely affected. These results demonstrate the need for *Rac1* in embryonic PDGFRα^+^ mesenchymal cells to generate the apical skull and dermis.

### *Rac1*-KO blocks the expansion of apical mesenchyme into meningeal and dermal layers

In normal development of the apical head, a thin layer of mesenchyme covers the brain at E12.5, called early migrating mesenchyme (EMM) (Dasgupta and Jeong 2019). This tissue does not directly contribute to the ossification process that forms the calvaria (Vu et al. 2021), but it primarily gives rise to meninges and dermis of the scalp. Osteogenic cells for calvaria originate from a different mesenchymal population located in a basolateral position near the eyes, called supra-orbital mesenchyme (SOM) (Dasgupta and Jeong 2019). From E13.5-E16.5, the SOM cells differentiate into bone primordia while migrating apically through the EMM as it undergoes differentiation into dermis and meninges. Thus, a multilayered structure with two different mesenchymal sources develops into SOM-derived calvaria between EMM-derived meninges and dermis (Dasgupta et al. 2019; Dasgupta and Jeong 2019; Ferguson and Atit 2019). The calcium-binding protein S100a6 marks mesenchymal cells of the pial and arachnoid layers of the meninges at E14.5 and later (DeSisto et al. 2020). At E12.5-E13.5, before the entrance of SOM-derived osteogenic cells, we found S100a6 to be broadly expressed in EMM of the *Rac1*-Wt apical head, colocalizing with Tomato from *Pdgfra-Cre* lineage tracing (Fig. 3A-B). The contractile protein αSMA has been used as a marker for mechanosensitive fibroblasts of the apical head, which specifically appear between E12.5-E14.5 and later give rise to the scalp dermis (Angelozzi et al. 2022; Tsujikawa et al. 2022). At E12.5, we found αSMA expression overlapping with S100a6 in the EMM. At E13.5-14.5, αSMA became restricted to the upper half of the S100a6^+^ mesenchyme, suggestive of a developing dermis (Fig. 3A-C). Therefore, from E12.5-E14.5 before the arrival of osteogenic cells, the EMM expands from a simple layer of S100a6^+^aSMA^+^ cells into a multilayered tissue composed of presumptive meninges (S100a6^+^) and dermis (S100a6^+^aSMA^+^) (Fig. 3D).

**Figure 3:**
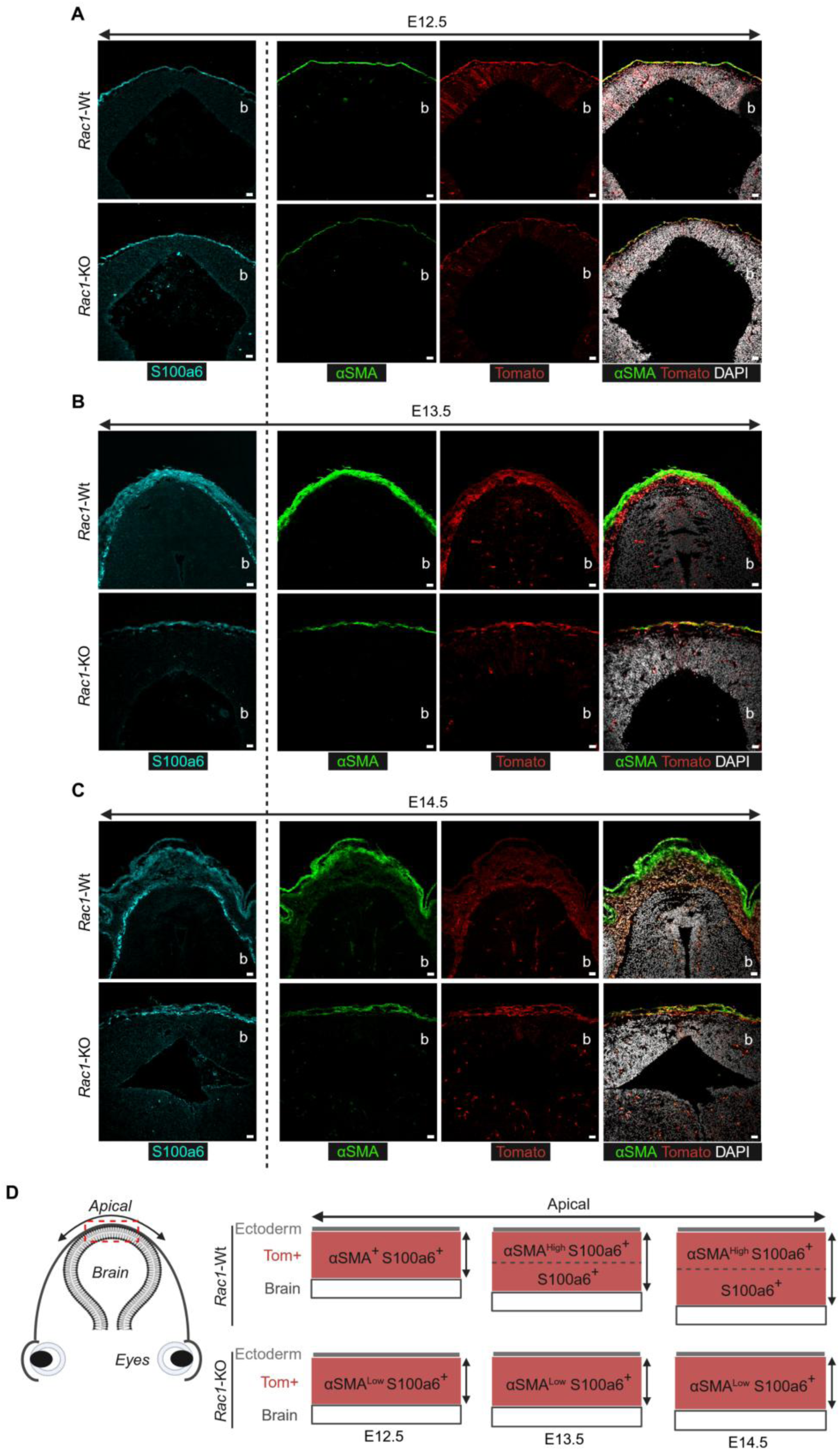
Deletion of Rac1 impedes the expansion of apical mesenchyme to form a multilayered tissue with αSMA^+^ dermal layer. (A-C) Immunostaining of S100a6, αSMA, and DAPI for E12.5-E14.5 *Rac1*-Wt and -KO (n=3). b=brain. Scale=50μm. (D) Graphical representation of apical head development. From E12.5-E14.5, the Tomato^+^ apical mesenchyme expands from a simple layer of αSMA^+^S100a6^+^ cells into a multilayered tissue with αSMA^+^S100a6^+^ dermal cells overlying αSMA^neg^S100a6^+^ meningeal cells. In the mutant αSMA is low and layers do not expand.

To explore the origin of the *Rac1*-KO phenotype, we examined these markers from E12.5-E14.5. The Tomato^+^ EMM in *Rac1*-KO was like *Rac1*-Wt at E12.5, but it failed to expand from E13.5-E14.5 (Fig. 3A-C). S100a6 was detected in *Rac1*-KO EMM at all time points, suggesting the maintenance of meningeal development. However, αSMA was noticeably weaker in *Rac1*-KO at E12.5 compared to *Rac1*-Wt, which is the earliest defect we observed. αSMA remained very low in the thin, discontinuous apical mesenchyme of *Rac1*-KO at E13.5-E14.5 (Fig. 3A-C). Reduced αSMA in the E12.5-E14.5 mesenchyme suggests an early failure of *Rac1*-KO cells to respond to the mechanical environment of the apical head.

At E18.5, S100a6 expression continued in the leptomeninges of *Rac1*-Wt as expected (DeSisto et al. 2020), and S100a6 was unchanged in *Rac1*-KO, which lacked calvarium (Suppl. Fig. 3C). This suggests maintenance of meningeal development in *Rac1*-KO, at least in part, but this needs more investigation. On the other hand, αSMA was not expressed in the apical dermis at E18.5 in either genotype (Suppl. Fig. 3C). This indicates that the αSMA^+^ cell state is transient from E12.5-E14.5, potentially due to forces present in the apical mesenchyme before the arrival of osteoprogenitors (Tsujikawa et al. 2022). αSMA was not detected in the basolateral mesenchyme of either genotype from E12.5-E14.5 (Suppl. Fig. 4A), suggesting restriction of the mechanosensitive cell state to the apical mesenchyme.

### Apical expansion of calvaria progenitors is defective in *Rac1*-KO

As described above, calvaria bones arise from mesenchymal progenitors in the SOM, which are established in a basolateral position above the eyes by E10.0 – E12.5. These osteoprogenitors form the calvaria bone primordia that migrate apically with the onset of mineralization at E14.0 – E15.0 and eventually meet to form sutures (Yoshida et al. 2008; Deckelbaum et al. 2012; Ferguson and Atit 2019). To examine whether there were defects in this population, we examined alkaline phosphatase (AP) activity, a pre-osteogenic marker, at E13.5 (Suppl. Fig. 4B). There was a significant reduction in the AP area (Suppl. Fig. 4B) within the basolateral mesenchyme in *Rac1*-KO, indicating a decrease in calvaria osteoprogenitors. This is consistent with basolateral hypoplasia at E14.5 in *Rac1*-KO (Fig. 2C inset 3) and with severe calvaria defects before birth (Fig. 1D). These results suggest that deletion of *Rac1* from PDGFRα^+^ cranial mesenchyme affects the development of AP^+^ osteoprogenitors in the SOM. This is in agreement with previous analysis of *Osx-Cre;Rac1^Flox/Flox^* preosteoblasts with defective differentiation (Huck et al. 2020).

### Rac1 regulates apical mesenchyme proliferation

A functional relationship between mechanical force from the growing brain, αSMA expression, and proliferation of the apical mesenchyme has been proposed (Tsujikawa et al. 2022). To determine whether hypoplastic apical head mesenchyme involves a lack of cell proliferation, 5-ethynyl-2’-deoxyuridine (Edu) staining was performed, and double-positive (Tom^+^Edu^+^) cells were quantified. *Rac1*-KO apical mesenchyme exhibited a significant reduction in double-positive cells at E12.5-E14.5 (Fig. 4A). Mesenchyme in the basolateral region exhibited the same trend but with significance only at E13.5-E14.5 (Fig. 4A). Previous studies have shown the need for Rac1 in G2/M progression and proliferation of fibroblasts and other mesenchymal cells (Moore et al. 1997; Nikolova et al. 2007; Gahankari et al. 2021). *In vitro* culture of *Rac1*-Wt and -KO dermal fibroblasts demonstrated reduced proliferation of *Rac1*-KO fibroblasts (Fig. 4B). To investigate whether *Rac1* is involved in fibroblast mechanosensing, we examined *Rac1*-Wt and -KO fibroblasts plated on tissue culture plastic, as the rigid surface causes cells to adopt a myofibroblast-like phenotype with αSMA and F-actin stress fibers (Son et al. 2024). *Rac1*-KO fibroblasts did not adopt the flattened morphology of Wt cells, and they expressed minimal αSMA, but they did form F-actin stress fibers with cortical actin (Fig. 4C). This experiment agrees with previous work on *Rac*1-KO MEF cells (Guo et al. 2006) and shows that *Rac1*-KO fibroblasts have deficient mechano-sensing ability, resulting in low αSMA expression and reduced proliferation. Taken together, these results suggest that apical fibroblasts require Rac1 to respond to mechanical stress produced by the growing brain in this specific region. Rac1 is needed for αSMA expression and for cell proliferation that expands the EMM into a population of dermal progenitor cells. Previous studies have shown the involvement of Rac1 in cellular mechanics (ruffling/lamellipodium/exertion of forces) inducing motility, proliferation, and deformation of mesenchymal cells by mechano-sensing of the microenvironment (Pasapera et al. 2015; Kunschmann et al. 2019; Marston et al. 2019).

**Figure 4:**
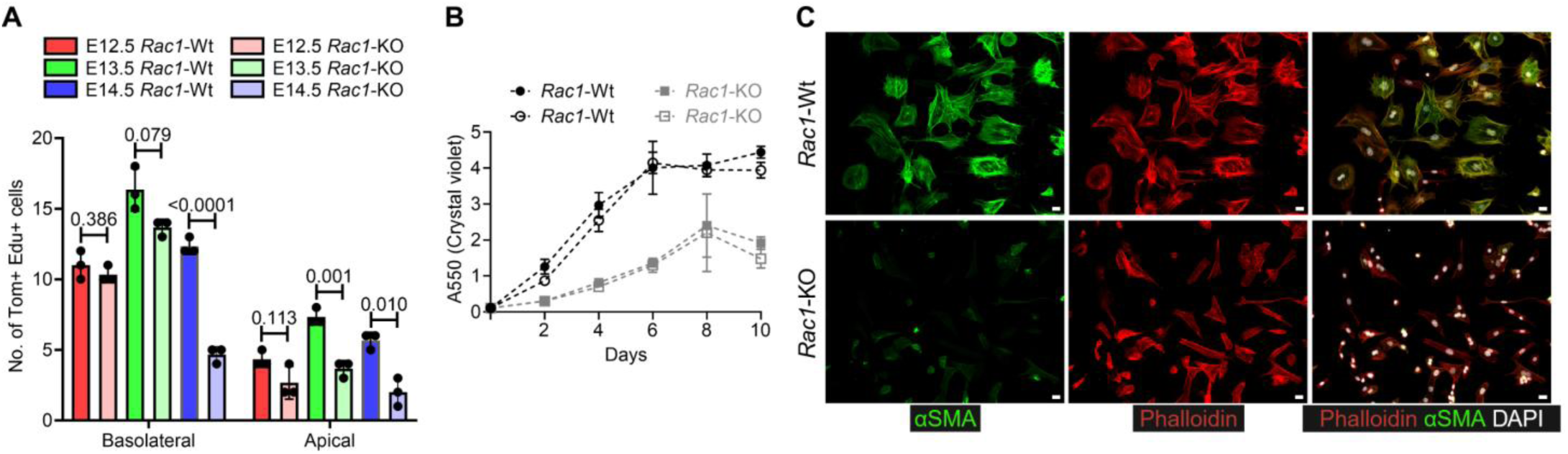
Loss of Rac1 attenuates the proliferation of apical mesenchyme. (A) Quantification Tom^+^Edu^+^ cells in basolateral and apical mesenchyme for E12.5-E14.5 (n=3), Data represented as mean ± SD, Unpaired t-test with Welch’s correction. (B) *In vitro* proliferation (Crystal violet assay) for *Rac1*-Wt and -KO dermal fibroblasts (n=2 biological replicates per genotype with 3 technical replicates), Data represented as mean ± SD. (C) Immunostaining of Phalloidin and αSMA for *Rac1*-Wt and -KO dermal fibroblasts (n=3) Scale=50μm.

### RAC1 regulates SRF expression and contractile function in fibroblasts

SRF is a ubiquitously expressed transcription factor that responds to serum and growth factors to promote gene expression programs governing cell proliferation and mechanotransduction. SRF achieves this by interacting with two co-factors: ternary complex factors (TCFs) and myocardin-related transcription factors (MRTFs) (Esnault et al. 2014; Gau and Roy 2018). SRF-TCF promotes immediate early gene expression and cell proliferation, while SRF-MRTF regulates actin dynamics, contraction, and cell motility (Esnault et al. 2014; Gualdrini et al. 2016; Gau and Roy 2018), including mechanotransduction signaling (Foster et al. 2017). The SRF protein is known to autoregulate its gene transcript, *Srf*, and its target genes, like *Acta2,* through upstream serum response elements (SRE) (Spencer and Misra 1996; Miano 2003; Deshpande et al. 2022). However, some mechanisms by which SRF is regulated remain obscure. A few studies have shown the possible involvement of Rho GTPases in regulating SRF (Hill et al. 1995; Liu et al. 2003; Miralles et al. 2003). We previously demonstrated that siRNA knockdown of *Rac1* reduces SRF nuclear localization in NIH/3T3 cells and dermal fibroblasts (Yao et al. 2022). Given this, we examined SRF expression in *Rac1*-KO fibroblasts and found that SRF nuclear localization and overall expression were reduced compared to *Rac1*-Wt (Fig. 5A-C). There was also a reduction of αSMA protein (Fig. 5B) and its mRNA, *Acta2* (Fig. 5C). To test the consequence of *Rac1* deletion on contractile function, we performed collagen contraction assays with *Rac1*-Wt and -KO fibroblasts seeded in an adherent collagen gel, with or without exposure to serum. In this setup, serum activates SRF-MRTF to induce contraction. Without serum, there was no contraction with either genotype (Fig. 5D). In the presence of serum, *Rac1*-Wt displayed a 20% reduction of gel area by 24 hours after release from the plastic surface, but *Rac1*-KO fibroblasts failed to contract the collagen gel (Fig. 5D). We performed immunostaining to examine SRF expression in the apical mesenchyme at E12.5-E13.5. SRF was strongly nuclear at E12.5 in both *Rac1*-Wt and -KO (Fig. 5E). SRF expression was maintained at E13.5 in the expanding apical mesenchyme of *Rac1*-Wt, but the mutant showed reduced expression in the thin and fragmented tissue (Fig. 5F). These results show that Rac1 is needed for full SRF expression and function in fibroblasts, possibly by affecting the autoregulation of SRF via subcellular localization of the protein. This is consistent with the model that mesenchymal *Rac1*-KO leads to scalp defects by impairing SRF functions, resulting in poor mechanosensing and insufficient proliferation.

**Figure 5:**
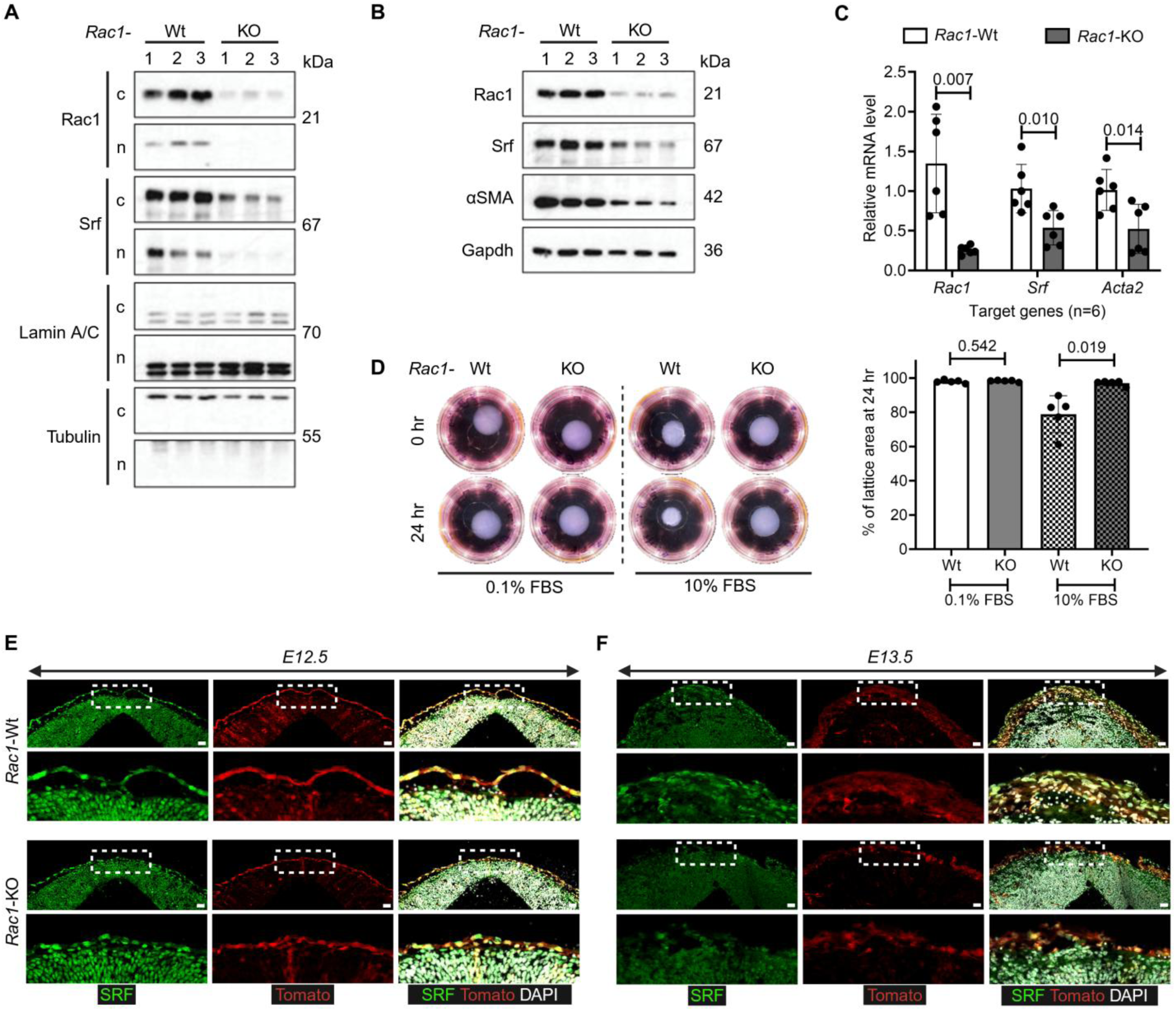
*Rac1*-KO fibroblasts exhibit reduced SRF expression and loss of contractile function. (A) Western blot with cytoplasmic (c) and nuclear (n) fractions (n=3). (B) Western blot for whole cell lysate showing reduction in SRF and its target gene αSMA (n=3). (C) qPCR data for *Rac1*, *Srf* and *Acta2* levels (n=6), Data represented as mean ± SD, Unpaired t-test with Welch’s correction. (D) Collagen lattice contraction, expressed as a percentage of the lattice area at 24hr with representative images at 0 and 24 hr after lattice detachment (n=5 biological replicates per genotype with 2 technical replicates). Data are plotted as mean ± SD, Unpaired t-test with Welch’s correction. (E-F) Immunostaining of SRF and DAPI for E12.5 and E13.5 *Rac1*-Wt and -KO (n=3). The inset represents the enlarged ROI shown below. Scale=50μm.

### Genetic deletion of *Srf* in the PDGFRα^+^ mesenchymal lineage causes apical head defects

To investigate the consequence of *Srf* deletion, we used *Srf^Flox/Flox^* mice (Miano et al. 2004) for the same conditional knockout strategy as was used for *Rac1*. Very few *Srf*-KO pups were born (2 mutants out of 69 pups, 17.25 mutants expected) and they displayed similar phenotype to that of *Rac1*-KO with defective development of the apical dermis and calvarium (Suppl. Fig. 5A). At E18.5, *Srf*-KO embryos were seen at ratios below the expected Mendelian ratio (9 mutants out of 114 pups, 28.5 mutants expected) (Fig. 6B). Those that were retrieved at E18.5 displayed misshapen apical heads (Fig. 6A,C) with significant reduction (>60%) in nasal, frontal, parietal, interparietal and occipital bones, compared to *Srf*-Wt (Fig. 6C, D). On the other hand, the skeleton of E18.5 *Srf*-KO displayed no additional defects other than reduced size (Suppl. Fig. 5C), which was like the *Rac1*-KO (Suppl. Fig. 1C). Immunostaining of SRF confirmed its absence in the E14.5 *Srf*-KO cranial mesenchyme (Suppl. Fig. 6A).

**Figure 6:**
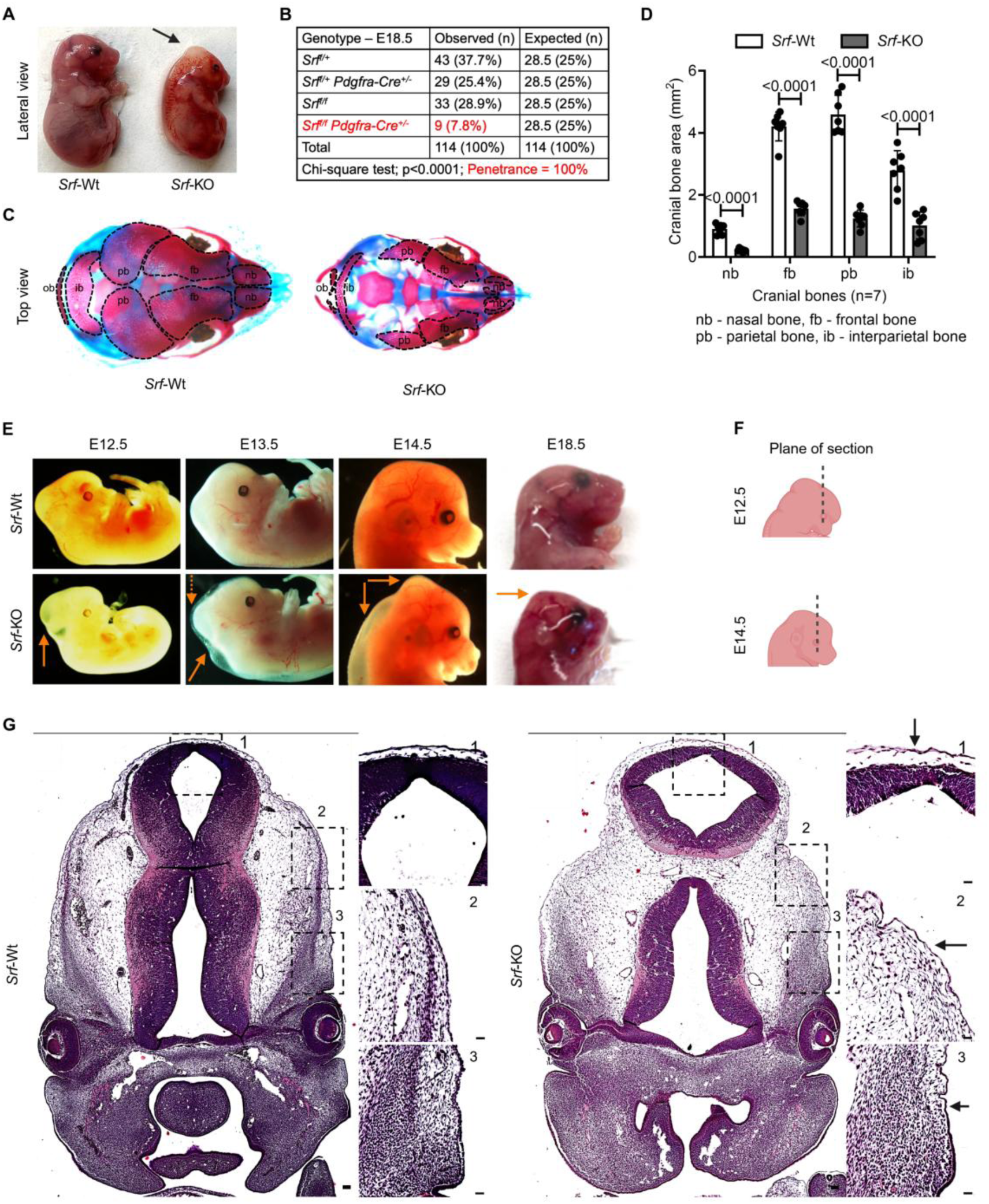
Mesenchymal *Srf*-KO causes scalp and calvaria defects and perinatal lethality like *Rac1*-KO. (A) *Srf*-Wt and -KO E18.5 (Lateral view) pups. The arrow indicates the apical head defect (n=9/9). (B) Genotypes of pups at E18.5, Chi-square analysis of genotypes with two-tailed P value, 100% of *Pdgfra^Cre^;Srf^Flox/Flox^* mice exhibit apical head defects. (C) Skeletal prep of the E18.5 skull with alizarin red and alcian blue staining (n=7). (D) Quantification of calvaria bones (n=7). Data represented as mean ± SD, Unpaired t-test with Welch’s correction. (E) Lateral view of *Srf*-Wt and -KO embryos from E12.5 to E18.5. The orange arrows indicate blebbing/edema, brain protrusion, and apical head defect (n=9-13). (F) Graphical representation of coronal plane of section for E12.5 and E14.5 mouse embryos shown in G and Suppl. Fig. 6B. (G) H&E staining of coronal sections of *Srf*-Wt and -KO E12.5 embryos. Dotted box=insets, arrow=hypoplasia of basolateral through intermediate and apical mesenchyme. The insets are represented as 1=apical, 2=intermediate, and 3=basolateral region of the head (n=1). Scale=50μm.

### Ontogeny of the *Srf*-KO cranial phenotype

To explore the origin of the phenotype in *Srf-*KO embryos, timed pregnancies were conducted. The earliest phenotype was observed at E12.5 with a slight change in the shape of the caudal head (Fig. 6E), followed by a protruding caudal head and a distinct blebbing at E13.5 (Fig. 6E). At E14.5, the caudal deformity enlarged with conical head shape and dorsal edema (Fig. 6E). Blebbing and edema were reduced by E18.5 (Fig. 6A), but the apical head defect remained externally obvious (Fig. 6A, E). A previous study targeting SRF in NCCs with *Wnt1-Cre* demonstrated fully penetrant facial clefting as early as E11.5 (Vasudevan and Soriano 2014). No such phenotype was observed in our *Srf*-KO, which indicates that despite the ability of *Pdgfra-Cre* to target NCCs (Soriano 1997; Tallquist and Soriano 2003; He and Soriano 2013; Mo et al. 2020) (Suppl. Fig. 2B-D), it acts later than *Wnt-Cre* and therefore gives low penetrance of facial cleft.

Coronal sections at E12.5 revealed hypoplasia of the basolateral pre-osteogenic region above the eye (Fig. 6G, inset 3) and continuing through the intermediate (Fig. 6G, inset 2) and apical mesenchyme (Fig. 6G, inset 1). Hypoplasia and edema persisted in these regions at E14.5 (Suppl. Fig. 6B). These data show the importance of SRF in PDGFRα^+^ mesenchymal cells that mediate apical head development and reveal a strikingly similar phenotype to *Rac1*-KO, consistent with the model that they work in the same pathway.

### *Srf*-KO affects the expansion of mechanosensitive fibroblasts during apical head development

To analyze the *Srf*-KO phenotype, we examined markers for the emergence of multilayered apical mesenchyme between E12.5-E14.5. Like *Rac1*-KO, S100a6 was detected in the hypoplastic *Srf*-KO apical mesenchyme at all time points (Fig. 7A-C, Fig. 3A-C). However, αSMA was severely reduced in the *Srf*-KO at all time points (Fig. 7A-C). Furthermore, the *Srf*-KO apical mesenchyme was thin and fragmented at E13.5-E14.5 like *Rac1*-KO. To determine whether hypoplastic apical head mesenchyme involves a lack of cell proliferation, Edu staining was performed, and Tom^+^Edu^+^ cells were quantified. Like *Rac1*-KO, the *Srf*-KO apical mesenchyme exhibited a significant reduction in double-positive cells at E12.5-E14.5 (Suppl. Fig. 7A). These results support our expectation that SRF is critical for cells in the apical mesenchyme to expand into αSMA^+^ dermal progenitors. Osteoprogenitors of the SOM were also affected by *Srf*-KO, with a significant reduction in alkaline phosphatase activity at E13.5 (Suppl. Fig. 7B). Mesenchyme in the basolateral region exhibited reduced proliferation (Suppl. Fig. 7A).

**Figure 7:**
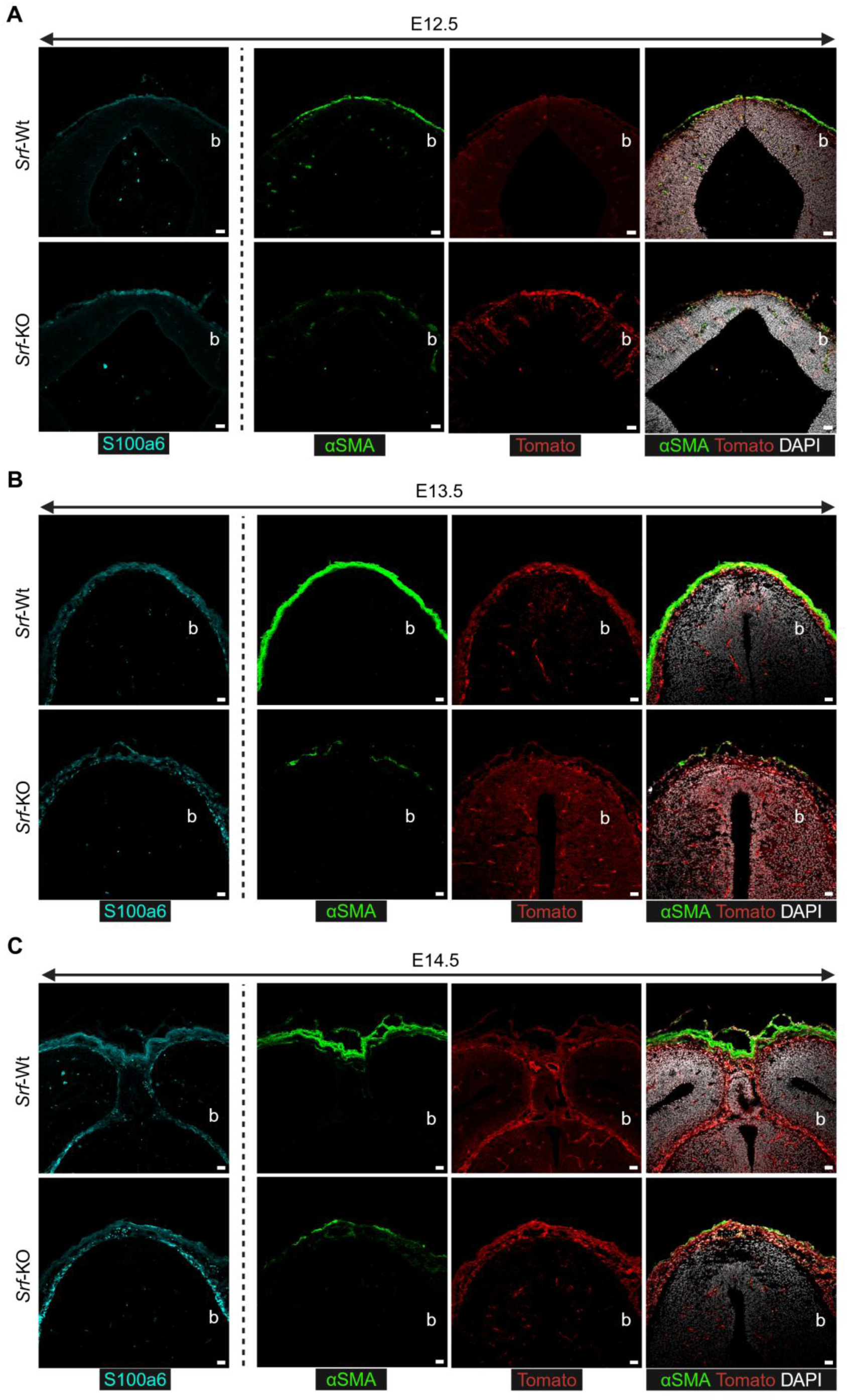
Mesenchymal deletion of *Srf* recapitulates the *Rac1*-KO apical mesenchyme phenotype with failure to expand the αSMA+ dermal layer. (A-C) Immunostaining of S100a6, αSMA, and DAPI for E12.5-E14.5 *Srf*-Wt and -KO (n=2-3). b=brain. Scale=50μm.

In conclusion, deletion of either *Rac1* or *Srf* in PDGFRα^+^ mesenchyme leads to aplasia of the scalp dermis and calvaria in a phenotype resembling severe AOS. Both Rac1 and SRF are critical for the expansion of scalp dermis, and for osteoprogenitors that form the calvaria bones. We propose that Rac1-SRF signaling operates in a mechanosensing pathway that is crucial for apical fibroblasts to expand into a population capable of creating scalp dermis. In AOS individuals, mutations in DOCK6 and ARHGAP31 are predicted to decrease Rac1 signaling, which could then lead to scalp and calvaria defects through the mechanisms described here.

## DISCUSSION

The scalp, calvaria, and meninges all originate from PDGFRα^+^ mesenchymal cells. A subset of these cells expresses contractile genes between E12.5-E14.5 (Angelozzi et al. 2022; Tsujikawa et al. 2022). The expression of αSMA (*Acta2*) in non-smooth muscle cells typically indicates mechanosensing properties, for example in myofibroblasts during wound healing (Hinz 2016). It was suggested that αSMA^+^ cells in the apical head are progenitors that expand into scalp dermal fibroblasts via mechanically regulated proliferation (Tsujikawa et al. 2022), but the significance of this population for apical head development has not been clarified. We found that knockout of *Rac1* caused downregulation of αSMA and reduced proliferation of PDGFRα+ mesenchymal cells. Consequently, there was aplasia of the scalp dermis and a phenotype at birth resembling AOS. *Srf* knockout led to a strikingly similar phenotype. Therefore, the key finding of our study is that αSMA^+^ progenitors of the apical head depend on Rac1 and SRF, and these cells are critical for scalp development. In both mutants, we also observed reduced proliferation and reduced alkaline phosphatase activity in the basolateral mesenchyme, which constitutes a distinct population of calvaria osteoprogenitors. Consequently, there was severe disruption of calvaria development with brain herniation, which occurs in severe cases of AOS.

We did not set out to study AOS. However, the calvaria phenotype associated with *Rac1*-KO intrigued us to investigate the development of the apical head. Mutations in ARHGAP31 and DOCK6, two regulators of RAC1/CDC42, have been linked to Adams-Oliver syndrome (AOS) as these mutations impair the regulators’ ability to render RAC1/CDC42 to its active form (GTP-bound). AOS individuals present defective dermis and bone at the vertex of the skull and truncated limbs at birth (Shaheen et al. 201 ; Southgate et al. 201), indicating faulty development of mesenchyme-derived tissues during development. Previous studies have demonstrated the role of *Rac1* in mesenchyme-derived tissues, but none of these studies reported any AOS-associated phenotypes (Fuchs et al. 2009; Suzuki et al. 2009; Thomas et al. 2010; Gahankari et al. 2021). On the other hand, our *Rac1*-KO recapitulated a phenotype similar to severe cases of AOS (Wehrens et al. 2020), with consistent scalp defects as observed in AOS (Shaheen et al. 2011; Southgate et al. 2011; Meester et al. 2015; Southgate et al. 2015). A genetic difference between AOS and our model is the loss of Rac1 itself, rather than mutation of Rac1 regulators, which may lead to a more penetrant and severe phenotype in our study.

Intriguingly, the *Rac1*-KO pups lacked limb defects/truncations seen in AOS newborns (Shaheen et al. 2011; Southgate et al. 2011). One reason could be that *Pdgfra-Cre^Tg^* was not expressed early enough to target limb bud mesenchyme. However, even with the expression of *Prrx1-Cre* at E9.5 (Logan et al. 2002), *Suzuki et al*. demonstrated short but not-truncated limbs in *Prrx1-Cre;Rac1^Flox/Flox^* mice (Suzuki et al. 2009), suggesting that loss of Rac1 in mesenchyme does not cause limb truncation irrespective of the timing of *Cre* expression. On the other hand, *Wu et al*. demonstrated various degrees of limb truncation in *Msx2-Cre;Rac1^Flox/Flox^* mice (Wu et al. 2008), which is similar to the limb defects observed in AOS individuals. It is important to note that *Msx2-Cre* targets limb bud ectoderm instead of mesenchyme, which suggests that the limb defect is due to loss of Rac1 in the ectoderm responsible for the patterning of limb mesenchyme. Thus, Rac1 appears to play different roles in limb and apical head development where different mechanisms govern tissue growth and patterning.

Previous studies demonstrated the significant role of Rac1 in mediating the cell cycle progression to G2/M transition and thereby proliferation (Moore et al. 1997). The loss of Rac1 or the expression of dominant negative Rac1 resulted in inhibition of DNA synthesis and cytostatic growth arrest (Moore et al. 1997; Assoian and Schwartz 2001; Nikolova et al. 2007). Rac1 regulates actin dynamics via its downstream effectors (WAVE, IQGAP, PAK and MLK2/3) to induce membrane ruffling/lamellipodia to accommodate cell growth, migration and cytoskeleton organization (Hall 1998; Ridley 2001; Aspenstrom et al. 2004; Bosco et al. 2009; Saci et al. 2011; Wojnacki et al. 2014). Expression of a dominant negative Rac1 (T17N) eliminated the “pioneer” leading edge microtubules of the cells, thereby limiting cell growth (Wittmann et al. 2003). These molecular mechanisms suggest some ways that Rac1 may operate in the expansion and maintenance of αSMA^+^ mechano-sensitive fibroblasts. Similarly, it is known that SRF regulates the expression of αSMA through SRF-MRTF signaling, which links cytoskeletal dynamics and gene expression (Spencer and Misra 1996; Schwartz 2010; Speight et al. 2016; Foster et al. 2017; Singh et al. 2023). In addition, SRF-TCF signaling promotes immediate early gene expression and cell proliferation in response to growth factors (Esnault et al. 2014; Gualdrini et al. 2016; Gau and Roy 2018). Although previous studies have suggested that Rho GTPases may regulate SRF in certain contexts (Hill et al. 1995; Liu et al. 2003; Miralles et al. 2003; Yao et al. 2022), it was unclear if Rac1-SRF axis was involved in apical head development. However, not only did our *Srf*-KO recapitulate the *Rac1*-KO phenotype but also provided molecular evidence of Rac1 signaling through SRF in the maintenance and expansion of mechanosensitive fibroblasts during apical head development. Thus, our study provides a new genetic insight into cranial development.

The *Pdgfra-Cre^Tg^* targeted the cephalic mesenchyme that gives rise to meninges, osteo and dermal progenitors. However, only the osteo and dermal progenitors were severely affected. The expansion and differentiation of meningeal progenitors still occurred, and we could still detect the expression of S100a6 at E18.5 for *Rac1*-KO. It may be that mechanosensitive fibroblasts do not contribute to meningeal development, but further genetic and lineage tracing studies are required to close this gap in our study.

Our goal was to understand the consequence of impaired Rac1 and SRF in apical head development, which incidentally resembled severe AOS scalp and calvaria defects. It is important to remember that the DOCK6 and ARHGAP31 mutations associated with AOS may affect CDC42 as well as Rac1. *Prrx1-Cre;Cdc42^Flox/Flox^* resulted in short limbs and body, abnormal calcification of the cranium, and cleft palate (Aizawa et al. 2012). Another study reported cleft lips and short snouts in mice using *Wnt-Cre;Cdc42^Flox/Flox^* (Liu et al. 2013). Therefore, future studies should investigate Cdc42 in mice with *Pdgfra-Cre*. In terms of SRF mechanisms, a previous study demonstrated heart outflow tract defects but no cranial defects when SRF was replaced with a mutant form that abrogated interaction with MRTF in neural crest mesenchyme. This suggests that SRF-MRTF interaction may be dispensable during cranial development (Dinsmore and Soriano 2022). Therefore, it would be interesting to know whether SRF-MRTF is required for apical head development by targeting their interaction with *Pdgfra-Cre*.

## EXPERIMENTAL MODEL AND SUBJECT DETAILS

### Mice

All animal experiments in this study were performed according to procedures approved by the Institutional Animal Care and Use Committee at the Oklahoma Medical Research Foundation. Mice were housed in groups of two to five and maintained in a 12-hour light/dark cycle with unlimited access to water and food. All strains were maintained on a mixed C57BL/129 genetic background at room temperature. Both males and females were analyzed with age-matched and littermate controls. Mouse lines *Pdgfra-Cre^Tg^ (JAX: 013148) (Berry and Rodeheffer 2013; Rivera-Gonzalez et al. 2016), ROSA26^Ai14^ (JAX: 007914) (Madisen et al. 2010), Rac1^Flox/Flox^ (Rac1^tm1Djk^, JAX: 005550) (Glogauer et al. 2003)* and *Srf^Flox/Flox^ (Srf^tm1Rmn^, JAX: 006658) (Miano et al. 2004)* were obtained from the Jackson Laboratories. For *time pregnancies*, females were checked for plugs every morning, and when plugged, the females were unset and considered as E0.5. For *in vivo* proliferation, 2 mM Edu was administered with 0.9% saline intraperitoneally to pregnant dams 1 hour prior to sacrifice.

### Primary cells

The dermal fibroblasts used in this study were isolated from E18.5 embryos (*Rac1^Flox/Flox^*) generated with *Pdgfra-Cre^Tg^*. Skin was dissected, floated dermis-side down on a sterile dish with 0.25% Trypsin and incubated at 37°C for an hour. The epidermis was removed, and the dermis was incubated with Collagenase II (500 U/mL) in Dulbecco’s Modified Eagle Medium (DMEM) at 37°C for an hour with intermittent trituration. The cells were filtered through a 100μm filter, followed by a wash with DMEM and the cells were plated in a growth medium (DMEM/F12, 10% FBS, 2mM L-Glutamine, 2mM Pen/Strep) incubated at 37°C. The rate of Cre-recombination was assessed by tomato reporter and 85-90% of the cells were tomato^+^. The experiments with primary cells were conducted at passage 2 and 3. For serum starvation experiments, the cells were kept in 0.1% FBS for 24 hours.

## METHOD DETAILS

### Western blotting

#### Whole cell extracts

The primary cells were lysed in ice-cold RIPA buffer (50 mM Tris pH-7.4, 150 mM NaCl, 1% NP-40, 0.1% SDS, and 0.25% Sodium deoxycholate) with protease and phosphatase inhibitors (Complete protease inhibitor cocktail, 1mM EDTA, 1mM Na_3_VO_4_, 1mM NaF, and 1mM PMSF). Post-10 minutes of incubation on ice, lysates were sonicated for 40 seconds. The lysates were then cleared by centrifugation to collect the protein supernatant, and the concentration was determined by BCA assay.

#### Cytoplasmic and nuclear extracts

The primary cells were resuspended in an ice-cold hypotonic buffer (20 mM Tris, pH-7.4, 10 mM NaCl, 3 mM MgCl_2_) with protease and phosphatase inhibitors (Complete protease inhibitor cocktail, 1mM EDTA, 1mM Na_3_VO_4_, 1mM NaF, and 1mM PMSF) to allow cell swelling. A 1/20 volume of NP-40 was added to each cell suspension, vortexed to disrupt the cytoplasmic membrane. The nuclei pellet was separated by centrifugation and the supernatant containing cytoplasmic extract was collected. The nuclei pellet was washed with a large volume of hypotonic buffer to remove cytoplasmic contamination, centrifuged and the pellet was resuspended in ice-cold nuclear extract buffer (10 mM Tris, pH-7.4, 100 mM NaCl, 1% Triton X-100, 0.1% SDS, 0.5% Sodium deoxycholate, 10% Glycerol and 1 mM EDTA) with protease and phosphatase inhibitors. The lysates were cleared by centrifugation and the protein concentration was measured by BCA assay.

The proteins were denatured with Laemmli sample buffer boiled at 95°C for 10 minutes and protein aliquots of 5-10 μg were separated parallelly by 8% or 12% denaturing gel (SDS-PAGE). The proteins were transferred onto a nitrocellulose membrane, washed with 1X TBST (1x TBS + 0.1%Tween 20), blocked with 5% BSA, and then subjected to detection with primary antibodies (1:2000 in 5% BSA) at 4°C O/N. The membranes were then washed thrice with 1X TBST and probed for secondary antibody (1:5000 in 5% milk) at RT for an hour, washed and developed with ECL substrate on autoradiography film.

### Whole-mount skeletal prep and microcomputed tomography

E18.5 embryos or P1 neonatal pups were processed for skeletal prep. Embryos/pups were de-skinned and fixed in 100% EtOH for 3 days followed by an Alcian blue stain (15 mg Alcian blue 8GX, 80 mL of 95% EtOH, and 20 mL of Glacial acetic acid) for 16 hours. The skeletons were then rinsed with 100% EtOH O/N, cleared in 1% KOH O/N, and counterstained with Alizarin red (5 mg Alizarin red S, 100 mL 1% KOH) for 8 hours. Skeletons were then cleared in 1%KOH followed by washes with gradient series of 1%KOH: glycerol and then stored in 100% glycerol. The cranial measurement was determined as an area of each pair of bones averaged and represented as a cranial bone area (mm^2^). For micro-computed tomography (micro-CT), the P1 pups were harvested and stored in 70% ethanol at 4°C and then imaged with a Viva CT 40 micro-CT (Scanco Medical) at an X-ray tube voltage of 70 kVp and an X-ray current of 114 µA as described elsewhere (Kwon et al. 2021).

### Histology and immunostaining of tissue

Embryos for paraffin processing were fixed for 24 hours at 4°C in Bouin’s fixative, rinsed under DI water, taken through an increasing ethanol gradient, cleared with xylene and finally infiltrated with parrafin without vacuum for an extended timeframe. For *histological stains*, paraffin blocks were sectioned (6 μm), baked at 56°C for 30 minutes and allowed to cool to room temperature. Sections deparaffinized in Histoclear and rehydrated by stepwise decreasing ethanol concentration to distilled water. For hematoxylin and eosin staining, slides were stained with Mayer’s hematoxylin for 1 minute and then washed with tap water. Slides were then incubated in Eosin Y for 20 seconds, washed with tap water, dehydrated, and coverslipped with permount. For *immunofluorescence*, embryos were fixed in 4% paraformaldehyde overnight at 4°C, washed with 1X PBS and preserved in 20% sucrose to make cryo blocks. The cryo blocks were sectioned (10 μm), washed, and blocked with 5% donkey serum in PBS for 1 hour, then incubated with primary antibody overnight at 4°C in a humidified chamber, then washed three times followed by incubation with fluorescent secondary antibody for 1 hour. The slides were washed and counterstained with DAPI. For *Edu detection*, the frozen sections were washed with 1X PBS, permeabilized, and incubated with Edu reaction cocktail (175 μL 1X PBS, 4 μL CuSO_4_, 0.2 μL 654 Azide dye, and 20 μL of 0.5 M Ascorbic acid) for 30 minutes in dark at room temperature. For co-staining, the sections were first immunolabelled and were then processed for Edu reaction. The sections were washed, counterstained with DAPI, and cover slipped with Flouro Gel. For *Alkaline phosphatase* staining, E13.5 embryos were fixed with 4% PFA for 30-40 minutes, washed with PBS, and transferred in 20% sucrose to dehydrate to prepare a cryo block. The sections were washed in PBST (1X PBS + 0.1% Tween 20) and TBST (1X TBS + 0.1% Tween 20) for 10 minutes each, followed by a 10-minute wash with NTMT (100 mM Tris, pH 9.4, 100 mM NaCl, 60 mM MgCl_2_ and 0.02% Tween 20). The sections were stained with 20µl/mL NBT/BCIP in the dark for 20 min at RT as described elsewhere (Ferguson et al. 2018). Slides were further washed in PBS and mounted with an aqueous mounting medium.

The sections were imaged on a slide scanner (Axiom), Nikon Eclipse 80i microscope, or Nikon AX-R confocal microscope.

### *In vitro* cell proliferation

Primary dermal fibroblasts of passage 2 were seeded on a 24-well plate at a density of 2.5 x 10^4^ cells/well (triplicates) in a growth medium (DMEM/F12, 10% FBS, 2mM L-Glutamine, 2mM Pen/Strep) incubated at 37°C. At an interval of 2 days, a plate was washed with 1X PBS, fixed in 10% NBF and washed with 1X PBS, then inverted to dry for 10 minutes. The plate was sealed with parafilm and stored at -20°C until all the time points were collected. Thaw the plate and stain with crystal violet (0.05% in dH_2_O) for 10 minutes, wash 3x with dH_2_O, and invert to drain well each time. Then extract with 1 mL of extraction buffer (10% Acetic acid) and measure at OD550.

### Collagen contraction assay

Primary dermal fibroblasts were cultured in three-dimensional type 1 collagen matrixes (collagen concentration, 1mg/mL; cell concentration, 1.0 x 10^6^ cells/mL). Matrixes were formed with 0.25 mL of cell/collagen solution placed on a pre-warmed 35 mm TPP dish and set for 5 minutes to polymerize. Upon initial polymerization, the matrixes were moved into an incubator for an hour to stabilize the gel matrix. The cell-collagen matrix was cultured in the growth medium (DMEM/F12, 10% FBS, 2mM L-Glutamine, 2mM Pen/Strep) incubated at 37°C for 24 hours, followed by 4 days with either 0.1% FBS serum starvation or with 10% FBS growth media. Mitomycin C (0.625 μg/mL) was added to the medium to suppress the effect of proliferation differences between controls and mutants. The matrix was replenished with fresh media every 48 hours. After 5 days of culture, the matrixes were digitally photographed before and after detaching gently from the dish and allowed to contract for 24 hours, photographed again as the final time point.

### RNA isolation and quantitative RT-PCR (qPCR)

Total RNA was isolated from primary dermal fibroblasts using Trizol. cDNA template was generated using PrimeScript RT enzyme, Oligo dT Primers, and random 6 mers. Quantitative PCR was performed with iQ SYBR Green master mix (Bio-Rad) on a CFX96 real-time PCR system (Bio-Rad). Bio-Rad CFX Manager (V2.1) software was used for analyzing cycle threshold (Ct) values and melting curves. Fold differences in transcript levels were normalized to the expression of *Timm17b*. Primer sequences are listed in Table 1.

### Statistical analysis

Data are presented as means +/− SD as indicated in the figure legends. Differences were analyzed by unpaired two-tailed t-test with Welch’s correction between two groups using Graphpad Prism 10. All measurements were from distinct biological samples (individual mice). Statistical parameters are found in the figure legends, including exact n and number of biological repeats. Each mouse was considered as a biological replicate.

## ACKNOWLEDGEMENTS

We thank Yuji Kondo and all members of the Olson laboratory for helpful discussions. We also thank the Microscopy Core Facility associated with P30-GM114731 of the Oklahoma Medical Research Foundation Centers of Biomedical Research Excellence. The Olson lab is supported by US National Institutes of Health (NIH) grants R01-AR073828 and R01-AR080896, the Oklahoma Center for Adult Stem Cell Research - a program of TSET, and grants from the Oklahoma City-based Presbyterian Health Foundation.

## AUTHOR CONTRIBUTIONS

**BHR-** Conceptualization, Methodology, Investigation, Visualization, and Writing - Original Draft, Review and Editing; **AR-** Investigation and Writing - Review and Editing; **HRK-** Investigation, Visualization, and Writing - Review and Editing; **WLB-** Investigation; **LEO-** Conceptualization, Funding Acquisition, Supervision, and Writing - Reviewing and Editing

## DECLARATION OF INTERESTS

The authors declare no competing interests.

---

**Supplemental Figure 1:**
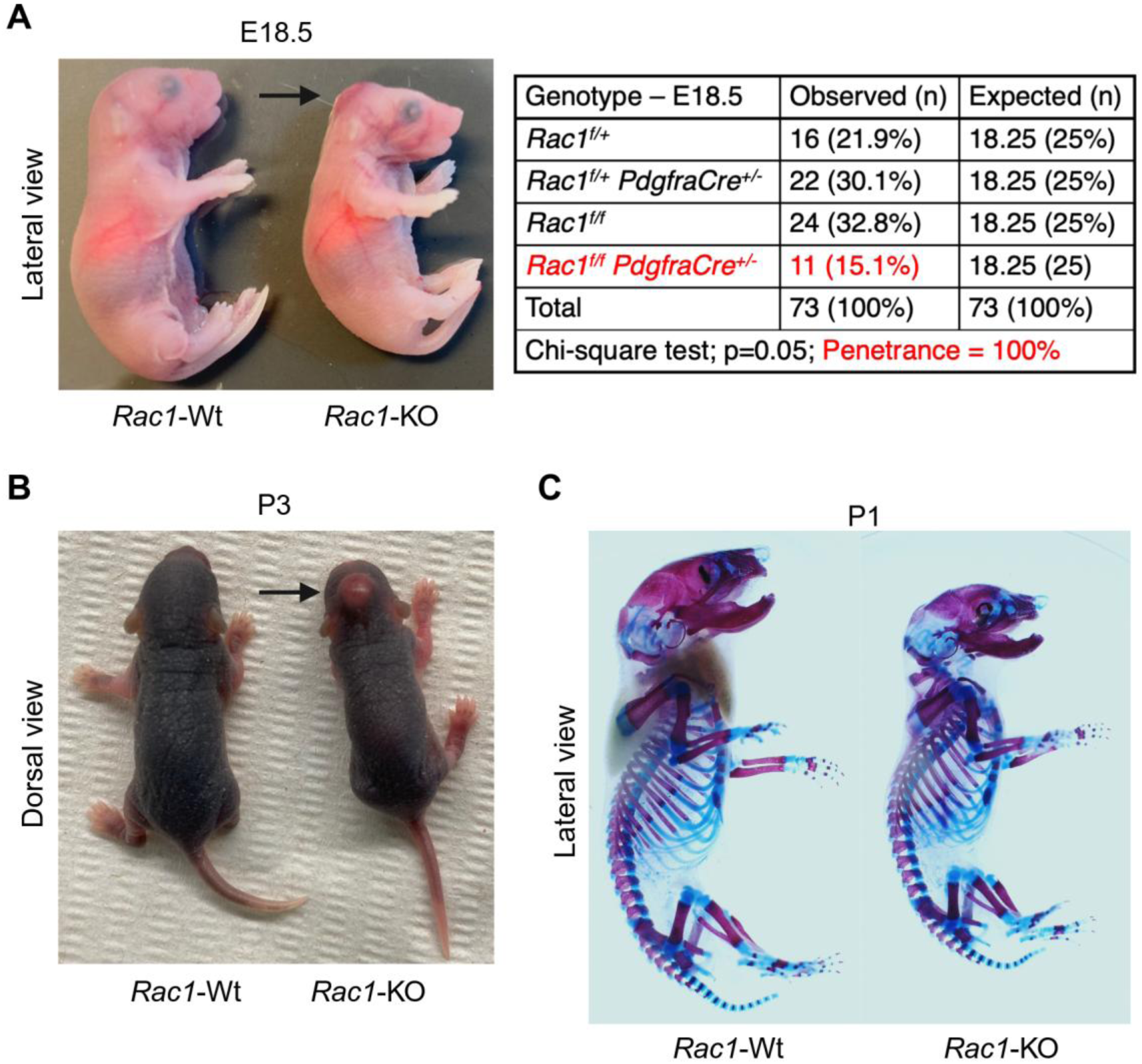
Deletion of mesenchymal *Rac1* causes scalp and calvaria defects but no limb truncation. (A) Lateral view of *Rac1*-Wt and -KO E18.5 pups. Arrow indicates the cranial defect (n=11/11). Genotypes of pups at E18.5, two-tailed Chi-square test of observed genotypes. (B) Dorsal view of *Rac1*-Wt and -KO P3 pups. Arrow indicates the cranial defect. (C) Skeletal preparation of the P1 pups with alizarin red and alcian blue staining (n=5).

**Supplemental Figure. 2:**
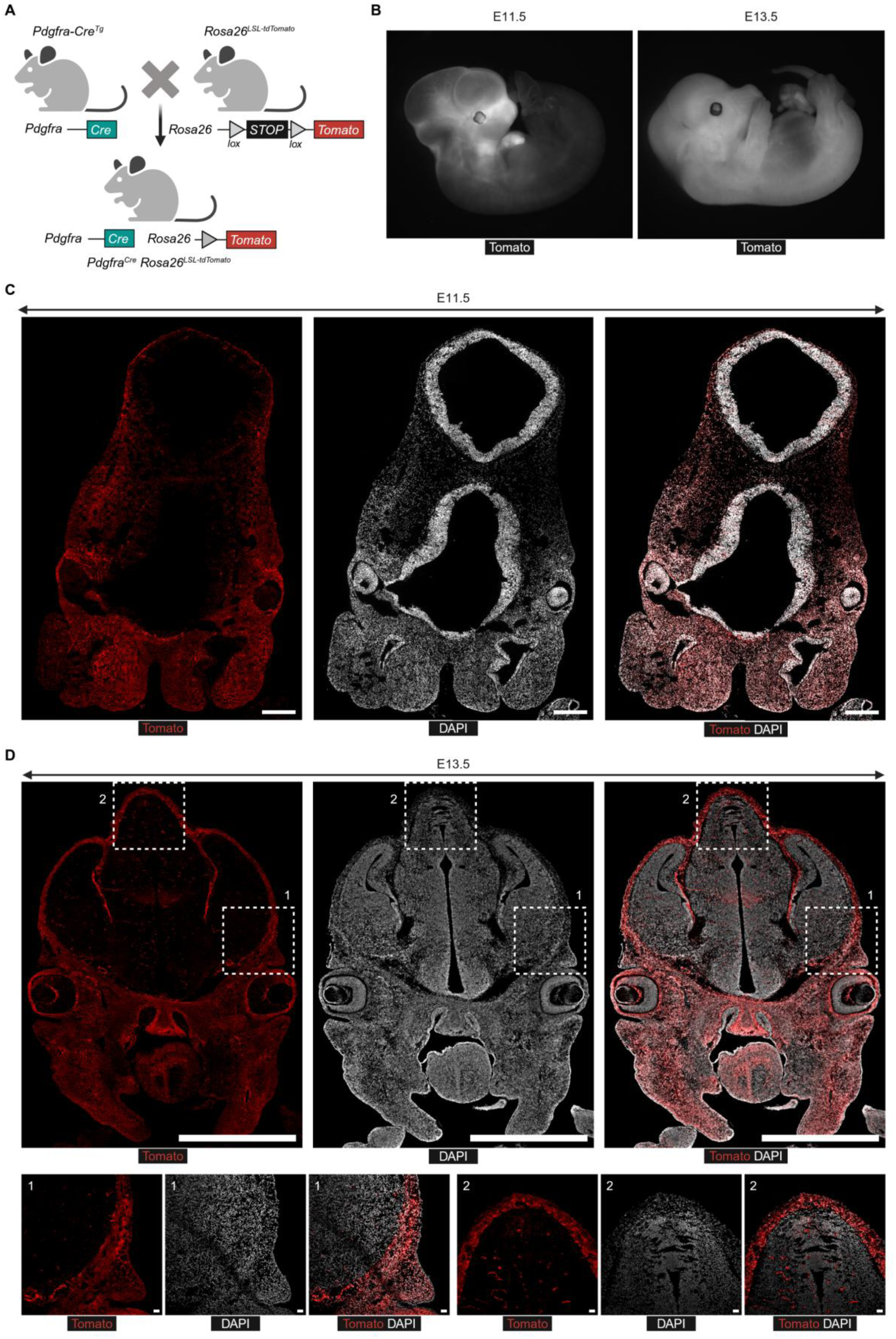
*Pdgfra-Cre^Tg^* activity in mesenchymal cells of mouse embryos. _(A)_ Graphical representation of genetic labeling strategy for *Pdgfra^Cre^ Rosa26^Ai14^*. (B) Tomato expression in E11.5 and E13.5 mouse embryos. (C-D) Coronal section of an E11.5 and E13.5 mouse embryo showing mesenchymal cells labeled with Tomato reporter and DAPI. The bottom panel represents the insets: 1=Basolateral and 2=Apical mesenchyme (n=2). Scale=100, 400, and 50μm.

**Supplemental Figure 3:**
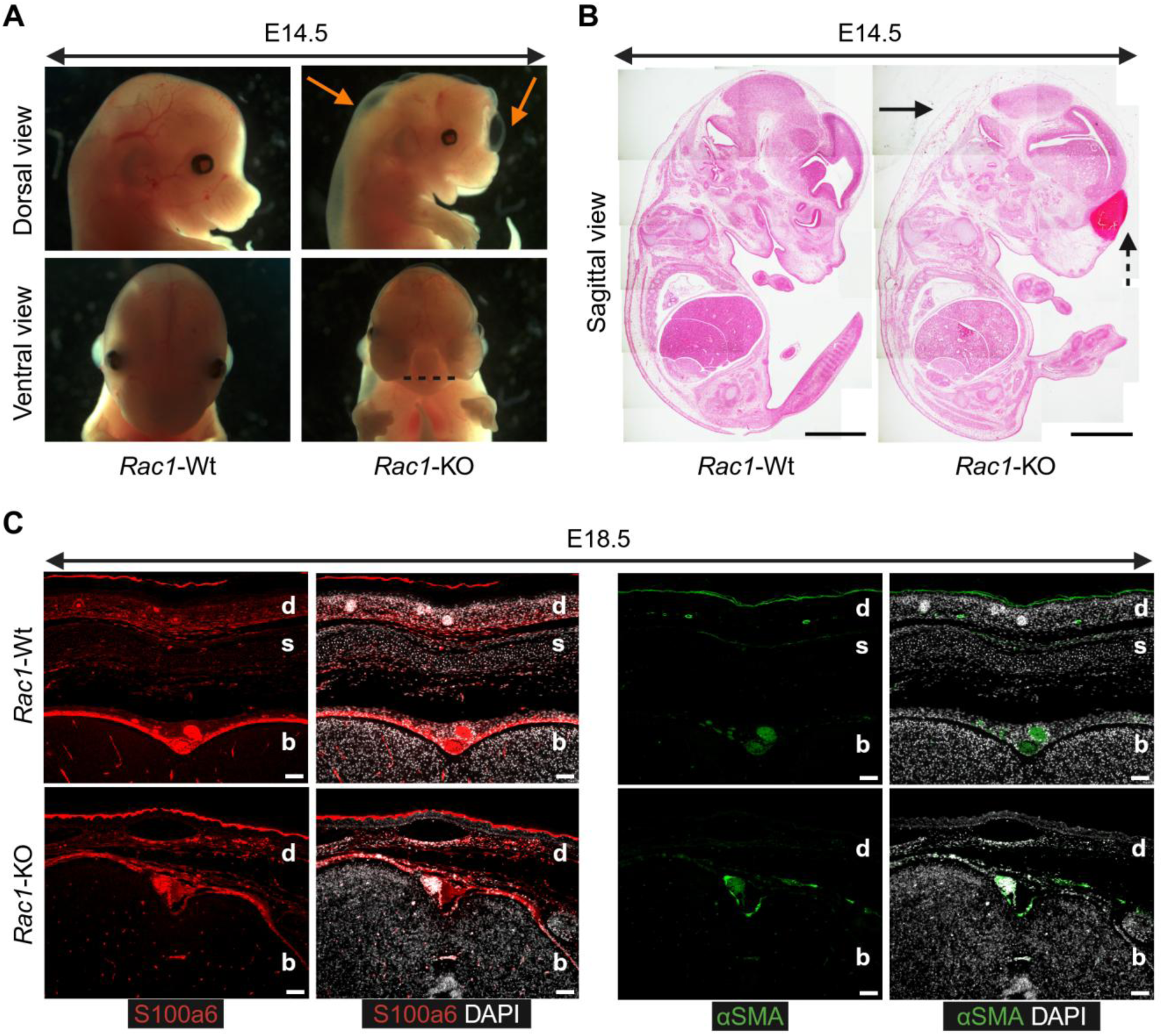
Mesenchymal *Rac1*-KO causes edema and dermal hypoplasia. (A) Dorsal and ventral view of *Rac1*-Wt and -KO embryos from E14.5. The orange arrows indicate blebbing/edema seen in all mutant embryos. The black dotted line indicates mid-facial cleft seen in 1 of 16 mutant embryos. (B) H&E staining of sagittal sections of *Rac1*-Wt and -KO E14.5 embryos (n=1). Arrow=edematous dorsal and apical mesenchyme, dotted arrow=hematoma. Scale=200μm. (C) Immunostaining of S100a6 and αSMA in the apical head at E18.5 for *Rac1*-Wt and -KO. S100a6 is expressed in leptomeninges at E18.5. αSMA is not expressed outside of the vasculature. d=dermis, s=skull and b=brain (n=1). Scale=100μm.

**Supplemental Figure 4:**
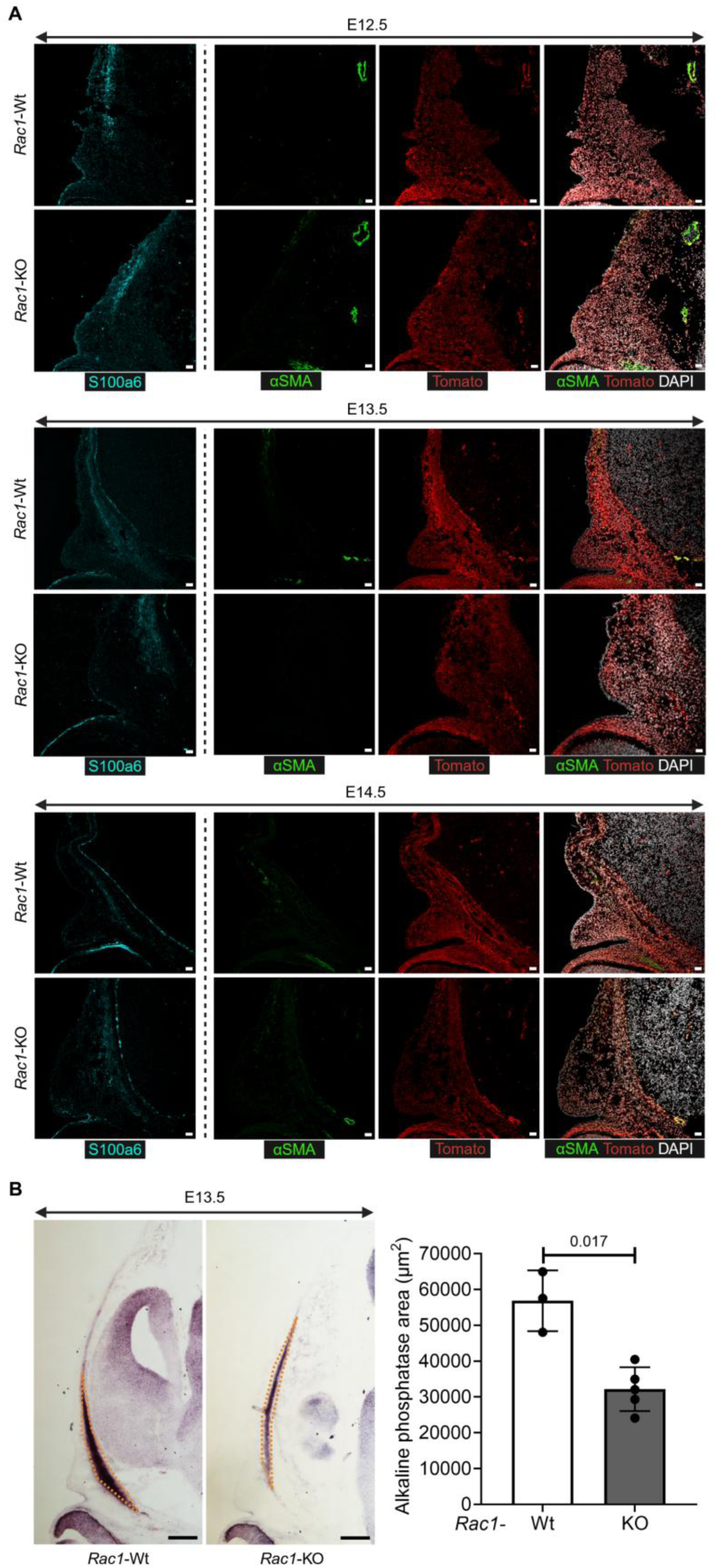
Mesenchymal *Rac1*-KO reduces osteogenesis. (A) Immunostaining of S100a6 and αSMA in the basolateral mesenchyme at E12.5-E14.5 for *Rac1*-Wt and -KO. Mesenchymal S100a6 is not affected in the mutant, and αSMA is not expressed outside of the vasculature. d=dermis, s=skull and b=brain (n=2-3). Scale=50μm. (B) Alkaline phosphatase staining for E13.5 *Rac1*-Wt and -KO (n=3). Scale=200μm. Dotted area=alkaline phosphatase area. Quantification of alkaline phosphatase area for E13.5 *Rac1*-Wt and -KO (n=3-5), Data represented as mean ± SD, Unpaired t-test with Welch’s correction.

**Supplemental Figure 5:**
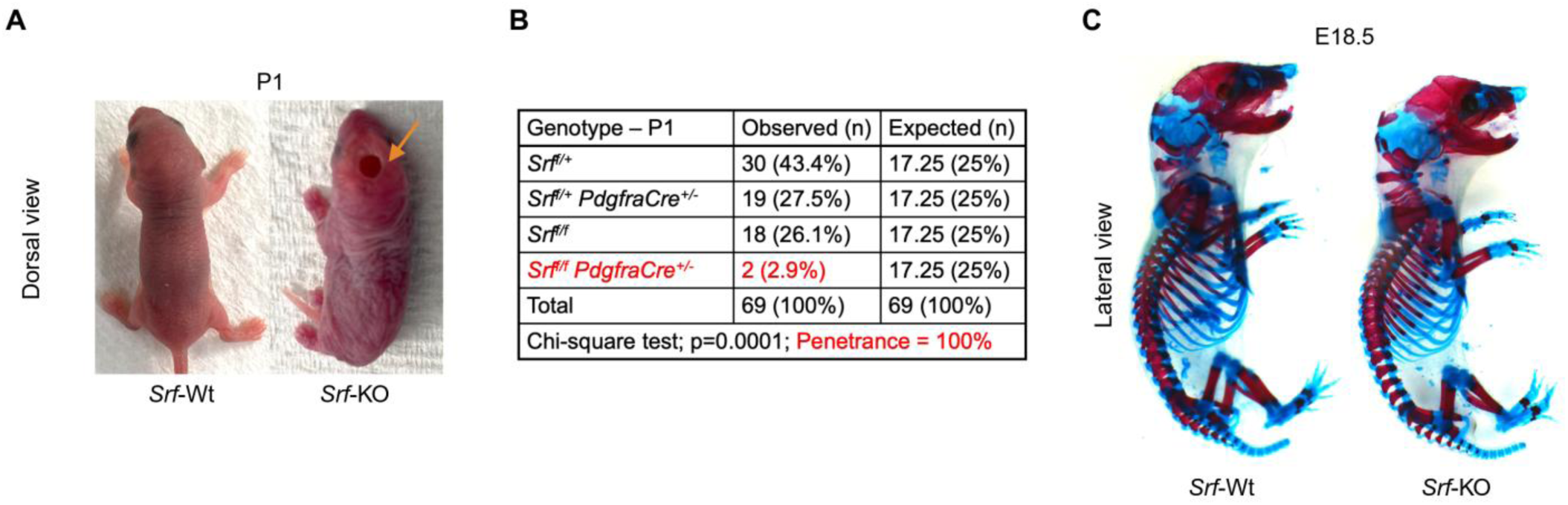
Deletion of mesenchymal *Srf* causes scalp and calvaria defects. (A) Dorsal view of *Srf*-Wt and -KO P1 pups. Arrow indicates the severe cranial defect with loss of calvaria and overlaying dermis. (B) Genotypes of pups at P1, two-tailed Chi-square test of observed genotypes. (C) Skeletal preparation of the E18.5 pups with alizarin red and alcian blue staining (n=7).

**Supplementary Figure 6:**
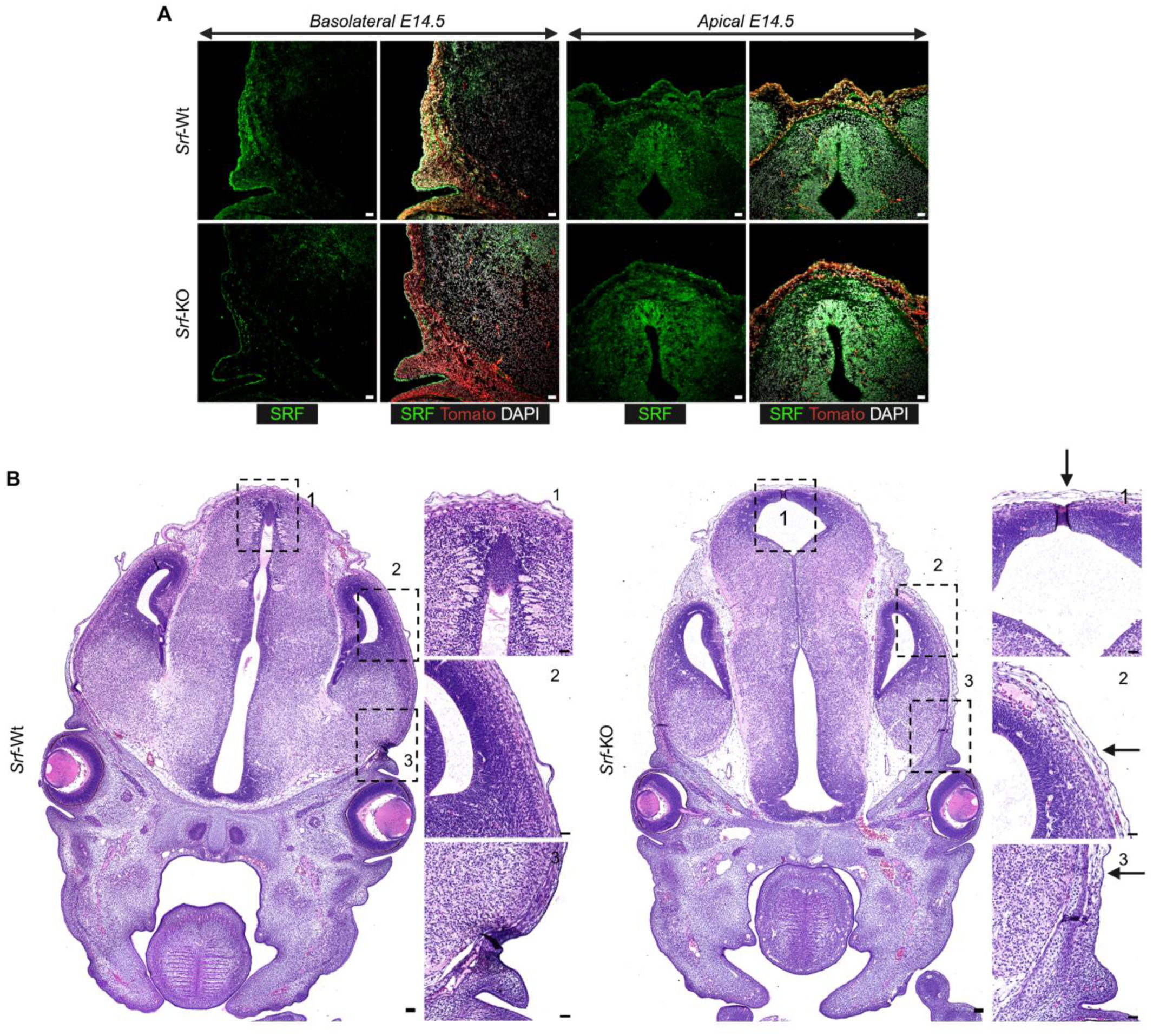
Deletion of *Srf* results in hypoplasia of basolateral and apical mesenchyme. (A) Immunostaining of SRF for *Srf*-Wt and -KO at E13.5 (n=1). Scale=100μm (B) H&E staining of coronal sections of *Srf*-Wt and -KO E14.5 embryos. Dotted box=insets, arrow=hypoplasia of basolateral through intermediate and apical mesenchyme. The insets are represented as 1=apical, 2=intermediate, and 3=basolateral region of the head (n=2). Scale=50μm.

**Supplemental Figure 7:**
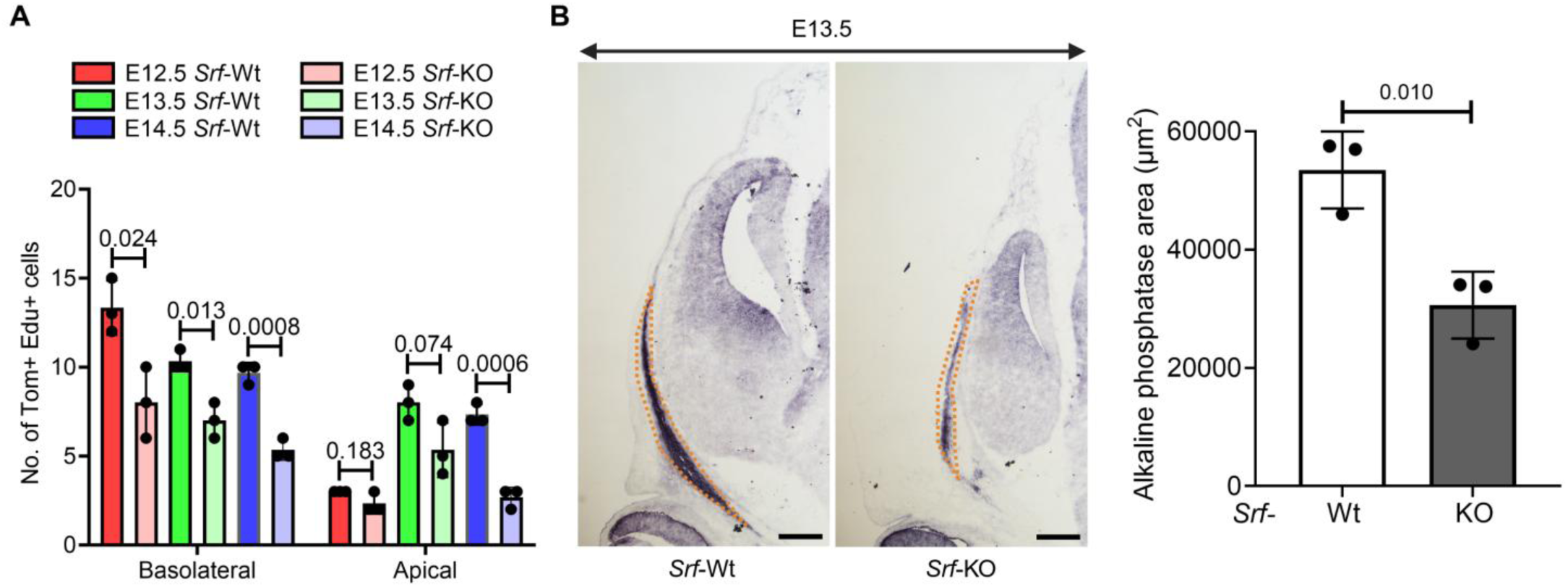
Mesenchymal *Srf*-KO reduces osteogenesis and attenuates proliferative cephalic mesenchyme. (A) Quantification Tom^+^Edu^+^ cells in basolateral and apical mesenchyme for E12.5-E14.5 (n=3), Data represented as mean ± SD, Unpaired t-test with Welch’s correction. (B) Alkaline phosphatase staining for E13.5 *Srf*-Wt and -KO (n=3). Scale=200μm. Dotted area=alkaline phosphatase area. Quantification of alkaline phosphatase area for E13.5 *Srf*-Wt and -KO (n=3), Data represented as mean ± SD, Unpaired t-test with Welch’s correction.

## Notes

### Competing Interest Statement

The authors have declared no competing interest.

